# Nuclear phylogenomics illuminates deep lineage divergence and complex evolutionary histories in Dipsacales

**DOI:** 10.64898/2026.11.14.766349

**Authors:** Dongzhu Jiang, Jiawei Yan, Diego F. Morales-Briones, Jacob B. Landis, Yong-Xing He, Pei-Feng Liu, Hong-Xin Wang, Jun Wen, Hua-Feng Wang

## Abstract

Relationships among the five subfamilies of Linnina within Dipsacales remain contentious. Extensive nuclear gene-tree discordance and cytonuclear discordance involving Linnaeoideae complicate reconstruction of their early evolutionary history. We analyzed whole-genome resequencing data from 88 Dipsacales species representing all nine subfamilies and eight outgroups. Using Dipsacales-specific nuclear reference locus sets, we recovered 3,023 orthologous nuclear loci. Nuclear analyses consistently placed (Morinoideae + Zabelioideae) sister to (Linnaeoideae + (Valerianoideae + Dipsacoideae)) across inference methods and character-coding schemes. Despite strong backbone support, extensive gene-tree discordance persisted at deep nodes. Node-level analyses suggested that incomplete lineage sorting (ILS) was a major source of discordance, while evidence for ancient gene flow varied among nodes. At the Linnaeoideae + (Valerianoideae + Dipsacoideae) node, topology frequencies approached ILS-only expectations, whereas internal branch lengths supported a model incorporating gene flow. At the Morinoideae + Zabelioideae node, multiple analyses provided more consistent evidence of ancient gene flow. ILS-only simulations recovered the observed plastid placement of Linnaeoideae in 8.63% of replicates, indicating that ILS could generate this topology; ancient gene flow, potentially involving plastid capture, remains a possible explanation for the cytonuclear discordance. Divergence-time estimates placed four major Linnina backbone divergences between 56.87 and 48.77 Ma. These findings suggest that early Linnina diversification involved successive lineage divergences and extensive ancestral polymorphism sorting, potentially accompanied by ancient gene exchange among some ancestral lineages.

**Graphical Abstract:** 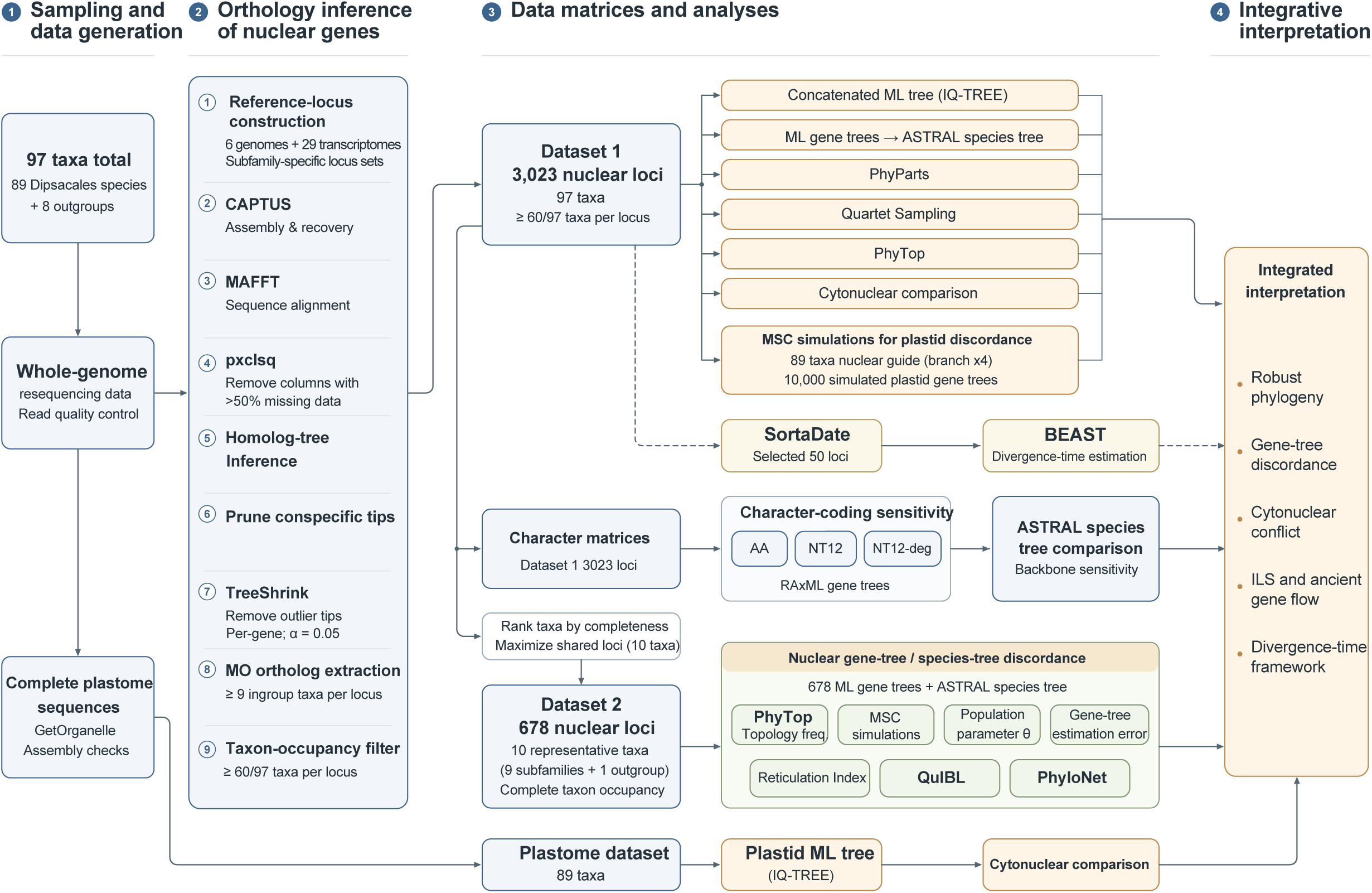

## 1 Introduction

Reconstructing the early evolutionary history of major plant lineages requires a well-resolved phylogeny and an understanding of the processes underlying lineage divergence. A species tree describes relationships among lineages, but individual gene trees may differ from this history. Ancestral polymorphisms can persist across successive speciation events when divergence intervals are short relative to ancestral coalescent times. Incomplete lineage sorting (ILS) can therefore produce discordance between gene trees and the species tree (Degnan and Rosenberg, 2006). Ancient gene flow can also contribute to discordance through the transfer of genetic variation among lineages (Yu et al., 2012). In addition, limited phylogenetic information at individual loci can lead to gene-tree estimation error (Roch and Warnow, 2015). Therefore, understanding early lineage divergence in plants requires more than a well-supported species tree (Cai et al., 2021; Morales-Briones et al., 2021). It also requires testing whether gene-tree discordance at different deep nodes is consistent with expectations under incomplete lineage sorting (ILS) and whether there is evidence for ancient gene flow.

Dipsacales contains anciently diverged lineages with diverse growth forms and reproductive structures, providing a suitable system for investigating these questions. The order comprises approximately 1,100 species, distributed mainly in temperate regions of the Northern Hemisphere (Bell and Donoghue, 2005a). Its extant members include both woody and herbaceous plants occurring in temperate forests, shrublands, grasslands, and alpine habitats. Some herbaceous lineages also extend into the South American Andes (Bell and Donoghue, 2005b). Floral and fruit morphology varies substantially across the order. Corollas show radial or bilateral symmetry, and stamen numbers differ among lineages. Fruits include drupes, berries, capsules, and achenes. Modifications of the calyx, epicalyx, and other fruit-associated structures contribute to fruit protection and dispersal (Donoghue et al., 2003; Carlson et al., 2009). This diversity provides a basis for examining the evolution of growth forms and reproductive traits across major lineages. Interpreting these differences and similarities requires a clear understanding of their phylogenetic relationships and divergence history.

Earlier morphological and molecular studies established two major clades within Dipsacales: Adoxaceae, comprising Opuloideae and Adoxoideae, and Caprifoliaceae sensu lato (Donoghue et al., 1992, 2001; Bell et al., 2001; Zhang et al., 2003). Subsequent plastid and nuclear phylogenomic studies supported Diervilloideae as sister to the remaining subfamilies of Caprifoliaceae. These studies also recovered Caprifolioideae as sister to Linnina (Wang et al., 2020; Xiang et al., 2020; Lee et al., 2021). The main unresolved relationships concern the branching order of the five Linnina subfamilies: Linnaeoideae, Morinoideae, Zabelioideae, Valerianoideae, and Dipsacoideae (Fig. S1). Within this clade, the positions of *Zabelia* and Linnaeoideae remain particularly contentious. Wang et al. (2021) used custom target-capture probes to obtain more than 700 nuclear exon loci for concatenation and coalescent-based analyses. Their concatenated RAxML analysis recovered (Morinoideae + Zabelioideae) as sister to (Linnaeoideae + (Valerianoideae + Dipsacoideae)). In contrast, their ASTRAL analysis did not recover Dipsacoideae as monophyletic. Instead, *Scabiosa* was recovered as sister to Valerianoideae, whereas *Dipsacus japonicus* was sister to Linnaeoideae (Wang et al., 2021). Lee et al. (2021) used Angiosperms353 target-capture data to construct two datasets comprising 313 exon-only loci and 308 exon-plus-flanking-region loci. Their exon-only ASTRAL analysis recovered the same five-subfamily backbone as the concatenation analysis of Wang et al. (2021). However, their exon-plus-flanking-region ASTRAL analysis and concatenated RAxML analyses placed *Zabelia* as sister to the other four subfamilies (Lee et al., 2021). Zuntini et al. (2024) subsequently inferred a large-scale angiosperm species tree using ASTRAL-MP and coding nucleotide sequences from the Angiosperms353 nuclear marker set. Its Dipsacales subtree recovered the same five-subfamily backbone as the concatenation analysis of Wang et al (2021). and the exon-only ASTRAL analysis of Lee et al (2021).

Beyond these differences among nuclear analyses, the placement of Linnaeoideae shows recurrent cytonuclear discordance. Wang et al. (2020) and Xiang et al. (2020) analysed datasets containing 68 and 75 plastomes, respectively. Both studies recovered Linnaeoideae as sister to (Morinoideae + Zabelioideae). This placement differs from the Linnaeoideae + (Valerianoideae + Dipsacoideae) relationship repeatedly recovered in nuclear analyses. Thus, two types of topological disagreement remain unresolved despite these advances. Nuclear datasets and inference methods differ in their recovery of *Zabelia* and Dipsacoideae monophyly. Nuclear and plastid data also support different sister-group relationships for Linnaeoideae.

Nuclear phylogenomic studies have also revealed extensive gene-tree discordance along the Linnina backbone (Wang et al., 2021; Lee et al., 2021). However, evidence for the role of ancient gene flow remains inconclusive. The nuclear phylogenetic network analyses of Wang et al. (2021) inferred candidate reticulation events involving Dipsacoideae, *Morina*, *Zabelia*, and lineages associated with Linnaeoideae. Their coalescent simulations addressed cytonuclear discordance and suggested that ILS alone could not explain some observed plastid relationships. These findings led the authors to propose ancient hybridization as an important source of phylogenetic discordance. By contrast, the SNAQ analyses of nuclear data by Lee et al. (2021) did not detect clear evidence of introgression. Nevertheless, the authors suggested that the history of Linnina might involve processes beyond ILS. They also noted that conserved single-copy loci could limit the detection of ancient gene flow.

Uncertainty in backbone relationships and the causes of discordance complicates interpretations of early lineage divergence and morphological evolution in Linnina. For example, the proposed sister relationship between *Zabelia* and Morinoideae informs whether their shared pollen characters can be interpreted as potential synapomorphies (Jacobs et al., 2011). The relative positions of Morinoideae and (Valerianoideae + Dipsacoideae) also underpin the hypothesis of independent origins of herbaceousness (Lee et al., 2021). Unresolved conflict mechanisms leave two further questions open: whether ILS primarily explains deep nuclear gene-tree discordance, and which ancestral lineages show evidence of ancient gene flow. Differences in taxon sampling, nuclear locus composition, and inference methods may affect both backbone reconstruction and the detection of ancient gene flow (Roch and Warnow, 2015; Shen et al., 2017; Lee et al., 2021). Broader nuclear locus sampling is therefore needed to assess the placements of *Zabelia* and Linnaeoideae and investigate discordance at the corresponding deep branches. This approach can clarify the branching order of major Linnina lineages and assess how well ILS explains nuclear gene-tree discordance. It can also identify ancestral branches associated with evidence of ancient gene flow.

To address these questions, we used whole-genome resequencing data from 97 taxa, including 89 Dipsacales species representing all nine subfamilies and eight outgroups. A Dipsacales-specific reference locus set was used to recover 3,023 orthologous nuclear loci. Nuclear and plastid data were combined to address two main questions : (1) which placements of Linnaeoideae and *Zabelia* receive robust support among competing nuclear topologies? (2) Why have previous studies reached different conclusions about ancient gene flow in Linnina, and how can the deep gene-tree discordance within this clade be explained? To investigate recurrent cytonuclear discordance, we assessed whether ILS could explain the plastid placement of Linnaeoideae relative to the nuclear species tree. Together, these analyses clarify relationships among major Dipsacales lineages and the processes underlying phylogenetic discordance, advancing our understanding of the order’s early evolutionary history.

## 2 Materials and methods

### 2.1 Taxon sampling

A total of 97 taxa were included in this study, comprising 89 Dipsacales species as the ingroup and eight non-Dipsacales species as outgroups. The eight outgroup taxa were *Columellia oblonga* (Columelliaceae), *Daucus carota* (Apiaceae), *Escallonia rubra* (Escalloniaceae), *Helianthus annuus* (Asteraceae), *Panax stipuleanatus* (Araliaceae), *Polyosma cambodiana* (Escalloniaceae), *Quintinia verdonii* (Paracryphiaceae), and *Ilex hylonoma* (Aquifoliaceae). We followed Lee et al. (2021) in recognizing two families within Dipsacales: Adoxaceae and Caprifoliaceae. Adoxaceae included Adoxoideae and Opuloideae. Caprifoliaceae included Caprifolioideae, Diervilloideae, Dipsacoideae, Linnaeoideae, Morinoideae, Valerianoideae, and Zabelioideae. The 89 Dipsacales species sampled in this study represented all nine subfamilies. For the species-rich Caprifolioideae and Opuloideae, taxon sampling was further refined to the section level. New sequencing data were generated from 53 field-collected samples. Leaf tissue from these samples was dried in silica gel. Data for the remaining 44 samples were obtained from previous projects and the NCBI database (Table S1).

### 2.2 Construction of Dipsacales-specific nuclear reference locus sets

A reference locus set for Dipsacales was constructed using publicly available data from six genomes and 29 transcriptomes across Campanulids with a major focus on Dipsacales (Table S2). Transcriptomes were processed following Morales-Briones et al. (2021) and assembled using Trinity v2.15.1 (Grabherr et al., 2011) with default settings, without read normalization. Open reading frames were predicted using TransDecoder v5.3.0 (Haas et al., 2013), with protein homology evidence from *Arabidopsis thaliana* and *Daucus carota*. Coding sequences were reduced with CD-HIT v.4.8.1 (-c 0.98 -n 10) (Fu et al., 2012). BLASTN 2.16.0+ (Camacho et al., 2009) searches were carried out with an *E-*value cutoff of ten and max_target_seqs set to 100. BLASTN 2.16.0+ hits were filtered with a minimal fraction of 0.4.

Homologs were clustered with MCL v.14-137 (van Dongen, 2000) with a minimum minus log-transformed *E* value cutoff of five, and an inflation value of 1.4, and only clusters with at least 17 taxa were retained. Individual homolog clusters were aligned using the OMM_MACSE v.12.01 pipeline (Scornavacca et al., 2019). Aligned columns were further cleaned to have no more than 70% missing data using ‘pxclsq’ from Phyx (Brown et al., 2017), and sequences with a gap ratio larger than 50% were identified with trimAl v.1.5 (Capella-Gutiérrez et al., 2009) and removed.

Homolog trees were inferred using IQ-TREE v.3.0.1 (Wong et al., 2026) using an extended model selection (Kalyaanamoorthy et al., 2017) and no clade support. Monophyletic and paraphyletic tips from the same species were pruned from the homolog trees, keeping only the tip with the highest number of characters in the trimmed alignment following Yang and Smith (Yang & Smith, 2014).

Long outlier tips were detected and removed using TreeShrink v.1.3.9 (Mai & Mirarab, 2018) with the ’per-gene’ mode and a false positive error rate threshold (α) of 0.05. Orthology inference was done using the tree-based ‘monophyletic outgroup’ (MO) approach from Yang and Smith (2014). The MO approach filters for clusters containing monophyletic, single-copy outgroup taxa. It then roots the tree by the outgroups, traverses the rooted tree from root to tip, and retains the branch with more taxa when gene duplication is detected. If no duplication is detected, we set the MO approach to output a one-to-one ortholog while also checking for the outgroup taxa being monophyletic and single-copy. Using the eight outgroups listed in Section 2.1, we retained orthologs containing at least nine taxa, yielding 7,085 orthologs, including 4,083 one-to-one orthologs.

Finally, for loci extraction, we split the ortholog FASTA files into the nine subfamilies of Dipsacales sampled for better loci recovery and accuracy: Valerianoideae (4168 loci) included samples from *Valeriana*; Linnaeoideae (6999 loci) included *Abelia*, *Dipelta*, *Kolkwitzia*, and Linnaea; Zabelioideae (6616) included *Zabelia*; Caprifolioideae (7077 loci) included *Heptacodium*, *Leycesteria*, *Lonicera*, and *Triosteum*; Adoxoideae (5838 loci) included *Adoxa* and *Sambucus*; Opuloideae (6216 loci) included *Viburnum*. Our transcriptome /genome (Table S2) sampling did not include any members of Morinoideae, so we included *Abelia*, *Dipelta*, *Kolkwitzia*, *Linnaea*, and *Zabelia* to create a reference with 7034 loci.

### 2.3 DNA extraction, read processing, and assembly

DNA was extracted from leaf tissue using a modified CTAB protocol (Doyle & Doyle, 1987). DNA quality and purity were assessed using a microspectrophotometer and 1% agarose gel electrophoresis. DNA concentration was measured using an Agilent 2100 BioAnalyzer. Samples containing at least 0.8 μg DNA were used for library preparation. Paired-end libraries were prepared with insert sizes of 300–500 bp and sequenced on the BGISEQ-500 platform at BGI (Shenzhen, China) (Mak et al., 2017). Raw reads were filtered using SOAPfilter v2.2 (Luo et al., 2012). Read pairs were removed when the proportion of ambiguous bases exceeded 10% of the read length or when low-quality bases (Phred ≤ 10) accounted for more than 40% of a read. Raw reads retrieved from NCBI were processed using the same workflow as the newly generated reads.

Nuclear loci were recovered using the captus_assembly workflow in CAPTUS v1.4.1 (Ortiz et al., 2026). Reads were further trimmed with the clean module (-trimq 20). Trimmed reads were assembled *de novo* with the assemble module (--min_count 5 --min_contig_len 300 --prune_level 3). The extract module used the subfamily-specific reference locus sets to recover nuclear sequences from the assemblies. Extraction thresholds were set to a Scipio score of 0.3, an identity of 85%, and a coverage of 50% (--nuc_min_score 0.3 --nuc_min_identity 85 --nuc_min_coverage 50). The align module collected the recovered sequences by locus for subsequent orthology inference.

Plastomes were assembled using GetOrganelle v1.7.7.1 (Jin et al., 2020) under default settings. Assembly graphs were inspected in Bandage v0.8.1 (Wick et al., 2015) to assess plastome completeness and inverted repeat (IR) boundaries. Plastomes were provisionally annotated using DOGMA (Wyman et al., 2004). Assemblies and annotations were then checked manually in Geneious R11.0.5 (Biomatters Ltd., Auckland, New Zealand).

### 2.4 Orthology inference and dataset construction

The overall workflow for orthology inference, dataset construction, and downstream phylogenomic analyses is summarized in Fig. S2. Nuclear orthologs were inferred using the tree-based workflow of Yang and Smith (2014), as implemented by Morales-Briones et al. (2022). Nuclear loci extracted with CAPTUS were aligned using MAFFT v7.520 (Katoh & Standley, 2013) with default settings. Alignment columns with >50% missing data were removed using pxclsq in Phyx (Brown et al., 2017). Within monophyletic or paraphyletic groups of conspecific sequences, we retained the sequence with the most non-ambiguous characters in the trimmed alignment (Yang & Smith, 2014). Anomalously long tips were removed using TreeShrink v1.3.9 in per-gene mode with α = 0.05 (Mai & Mirarab, 2018). Orthologs were extracted using the monophyletic-outgroup method (Yang & Smith, 2014). The eight taxa listed in Section 2.1 were designated as outgroups. All other taxa were treated as the ingroup. Each locus was required to contain at least nine ingroup taxa. Loci present in at least 60 of the 97 sampled taxa were then retained to form Dataset 1 (3,023 loci; Fig. S3). This dataset was used for nuclear phylogenetic inference, character-coding comparisons, and gene-tree discordance analyses. Dataset 2 was constructed from Dataset 1 to investigate the potential causes of discordance between nuclear gene trees and the species tree. A custom Python script was used to select one representative taxon from each of the nine subfamilies and one outgroup. Candidate taxa were ranked using a completeness score (S = 0.60P + 0.25E + 0.15[1 − M]), where P is locus occupancy, E is normalized total effective sequence length, and M is the proportion of missing characters. Up to five taxa with the highest scores were retained from each subfamily and the outgroup. All possible combinations of these candidates were evaluated. Combinations were ranked primarily by the number of orthologous loci shared by all 10 taxa. Ties were resolved using lower missing-data proportions and greater total effective sequence length. The highest-ranked combination included one representative from each subfamily and *Polyosma cambodiana* as the outgroup. The resulting Dataset 2 contained 678 orthologous loci with complete taxon occupancy (Table S3).

### 2.5 Phylogenetic analyses

To reconstruct the plastome phylogeny for comparison with the nuclear species tree, plastome sequences were aligned using MAFFT v7.520 (Katoh & Standley, 2013). Maximum-likelihood (ML) analysis was performed in IQ-TREE v3.0.1 (Wong et al., 2026) using a single partition. The best-fitting nucleotide substitution model was selected using ModelFinder (Kalyaanamoorthy et al., 2017). Branch support was assessed with 1,000 standard nonparametric bootstrap replicates (-b 1000).

To infer nuclear phylogenetic relationships and assess their consistency across inference methods, Dataset 1 (3,023 loci) was analyzed using concatenation and multispecies coalescent (MSC) approaches. For the concatenation analysis, locus alignments were combined usingPhyloSuite v1.2.3 (Zhang et al., 2020), with each locus treated as an initial partition. The best-fitting partitioning scheme and substitution models were selected in IQ-TREE v3.0.1 using ModelFinder with MFP+MERGE and relaxed clustering (-rcluster 10). An ML tree was inferred, and branch support was assessed with 1,000 ultrafast bootstrap replicates (-B 1000) and 1,000 SH-aLRT replicates (-alrt 1000). For the MSC analysis, ML gene trees were inferred separately for all 3,023 loci using IQ-TREE v3.0.1. The best-fitting substitution model was selected for each locus using ModelFinder, and branch support was assessed with 1,000 standard nonparametric bootstrap replicates (-b 1000). The best-scoring ML gene trees were used to infer a species tree using ASTRAL-III v5.7.8 (Zhang et al., 2018). Species-tree support was assessed using local posterior probabilities (LPP).

To assess the sensitivity of nuclear phylogenetic relationships to character coding, three character representations were generated from Dataset 1: amino acids (AA), first and second codon positions (NT12), and degeneracy-coded NT12 (NT12-deg). ML gene trees were inferred using RAxML v8.2.12 (Stamatakis, 2014) under the PROTGAMMALG model for AA and the GTRGAMMA model for NT12 and NT12-deg. Species trees were inferred using ASTRAL-III v5.7.8 and compared with the species tree inferred from the full nucleotide alignments of Dataset 1.

### 2.6 Phylogenetic discordance and cytonuclear conflict

Nuclear gene-tree concordance and discordance were quantified at each species-tree node using PhyParts (Smith et al., 2015). PhyTop was applied to the 3,023 gene trees and the nuclear species tree from Dataset 1 (Shang et al., 2025). This analysis assessed node-level signals of incomplete lineage sorting (ILS) and introgression/hybridization (IH). Quartet Sampling (QS) was used to assess branch support and discordance in the nuclear species tree (Pease et al., 2018). We performed 1,000 quartet replicates per internal branch, with gene partitions randomly sampled using the --genetrees option. Quartet topology likelihoods were calculated using RAxML-NG under the GTR+G4 model. The log-likelihood difference threshold between the best and second-best topologies was set to 2 (--lnlike 2). Quartet concordance (QC), quartet differential (QD), and quartet informativeness (QI) were then calculated.

Multispecies coalescent (MSC) simulations were used to test whether ILS alone could generate the relationships recovered in the empirical plastome tree. Dataset 1 was restricted to the same 89 taxa included in the plastome analysis. Species tree was inferred following the procedures described above and used as the simulation guide tree. Its branch lengths in coalescent units were multiplied by four, assuming a fourfold higher coalescent rate for plastid lineages than for nuclear lineages. We simulated 10,000 plastid gene trees using DendroPy and rooted them with *Ilex hylonoma* (Sukumaran and Holder, 2010). PhyParts was used to compare the simulated trees with the empirical plastome tree. We then calculated the frequency with which each relationship in the empirical plastome tree was recovered.

### 2.7 Sources of nuclear gene-tree discordance

Potential sources of nuclear gene-tree discordance across the nine-subfamily backbone were assessed using Dataset 2. The 678 maximum-likelihood gene trees and the corresponding ASTRAL species tree were analysed using PhyTop (Shang et al., 2025). Node-level signals of incomplete lineage sorting (ILS) and introgression/hybridization (IH) were assessed. Gene trees were simulated under the multispecies coalescent (MSC) model. Simulated and empirical discordance patterns were compared to test whether ILS alone could explain the observed conflict.

The population mutation parameter θ, node-reconstruction failure rates and a Reticulation Index were estimated following Cai et al. (2021). These metrics were used to further assess potential sources of discordance. θ was calculated from corresponding internal branch lengths in mutation and coalescent units. Gene-tree estimation error was assessed through parametric simulations on the fixed ASTRAL topology. Simulated sequences were matched to empirical locus lengths and substitution-model parameters. Node-reconstruction failure rates were defined as the proportions of simulated gene trees failing to recover the corresponding reference clades. Triplets were classified as unbalanced when their minor-topology frequency differences exceeded the maxima obtained for the corresponding triplets under MSC simulations. Candidate reticulation signals identified from these triplets were mapped onto the species tree. For each node, the Reticulation Index was calculated as the mapped signal count divided by the total number of associated triplets.

QuIBL was applied to Dataset 2 to compare ILS-only and ILS-plus-introgression models using internal branch-length distributions of rooted triplet gene trees (Edelman et al., 2019). All three-taxon combinations among the nine ingroup representatives were analyzed with *Polyosma cambodiana* as the outgroup. QuIBL was run with numdistributions = 2, likelihoodthresh = 0.01, numsteps = 40, and gradascentscalar = 0.5. We defined ΔBIC as BIC_ILS_ − BIC_ILS + introgression_. Discordant triplet topologies were classified as supporting ILS + IH (ΔBIC > 10), supporting ILS-only (ΔBIC < −10), or ambiguous (|ΔBIC| ≤ 10). Triplets were assigned to internal nodes according to the two descendant lineages and their nearest reference lineage in the species tree. For each node, the proportions of discordant-topology records classified as ILS-only, ILS + IH, or ambiguous were calculated.

### 2.8 Phylogenetic network inference

Phylogenetic network inference was performed on Dataset 2 using PhyloNet v3.8.2 (Than et al., 2008). Gene trees were rooted with *Polyosma cambodiana*, and the nine ingroup representatives were treated according to their corresponding subfamilies. Networks were inferred using the maximum pseudo-likelihood method implemented in InferNetwork_MPL (Yu & Nakhleh, 2015). The maximum number of reticulations (H) was varied from 0 to 5. For each maximum-reticulation setting, 200 searches were performed (-x 200), and the best-scoring network was retained (-n 1). For each value of H, the final total log pseudo-likelihood of the best network inferred by PhyloNet was compared. Following Blair and Ané (2020), H was selected by examining the change in the total log pseudolikelihood across increasing numbers of reticulations. The smallest H at the onset of the plateau was retained. Inheritance probabilities (γ) were estimated for the inferred reticulation edges.

### 2.9 Divergence time estimation

Divergence times were estimated using nuclear loci from Dataset 1. Loci were ranked with SortaDate (Smith et al., 2018) using default settings. Ranking was based on topological concordance with the Dataset 1 ASTRAL species tree, root-to-tip variance, and total tree length. The 50 highest-ranked loci were concatenated for dating analyses. Analyses were performed in BEAST v2.7.7 (Bouckaert et al., 2019) under a GTR + Γ4 substitution model with four gamma-rate categories, an uncorrelated lognormal relaxed clock (Drummond et al., 2006), and a Yule tree prior.

Six calibration points were specified as MRCA priors. Based on previous molecular dating estimates of the early divergence of Aquifoliales (Bremer et al., 2004, Wikström et al., 2015), we assigned a normally distributed secondary calibration prior to the Aquifoliales–Dipsacales divergence (mean = 100 Ma, σ = 6 Ma). The remaining five fossil calibrations were assigned lognormal priors. The *Heptacodium* fossil was used to calibrate the crown of Caprifolioideae, represented by the MRCA of *Heptacodium*, *Leycesteria*, *Lonicera*, *Symphoricarpos*, and *Triosteum* (offset = 20.4 Ma, M = 2.0, S = 1.25) (Popova et al., 2019). *Weigela weichangensis* seed was used to calibrate the MRCA of *W. floribunda* and *W. florida* (offset = 22.1 Ma, M = 2.0, S = 1.25) (Liang et al., 2013). *Lonicera* pollen was assigned to the MRCA of *Leycesteria*, *Lonicera*, *Symphoricarpos*, and *Triosteum* (offset = 34.0 Ma, M = 1.8, S = 1.25) (Wingate & Nichols, 2001). *Viburnum* pollen was used to calibrate the MRCA of *Sambucus* and *Viburnum* (offset = 47.0 Ma, M = 1.7, S = 1.25) (Donoghue, 1985). The fruit fossil *Patrinia hokiana* was used to calibrate the crown of Valerianoideae, represented by the MRCA of *Patrinia*, *Nardostachys*, *Valeriana*, *Valerianella*, and *Centranthus* (offset = 5.3 Ma, M = 2.2, S = 1.25) (Ozaki, 1980). All calibrated clades were constrained to be monophyletic. The MCMC analysis was run for 9×10^8^ generations, with parameters and trees sampled every 1,000 generations. Convergence was assessed in Tracer v1.7 (Rambaut et al., 2018), and effective sample sizes (ESS) >200 were required for all key parameters. The first 10% of samples were discarded as burn-in. A maximum clade credibility tree was summarized in TreeAnnotator using mean node heights, and 95% highest posterior density intervals (HPD) were reported for node ages.

## 3. RESULTS

### 3.1 Characteristics of the phylogenomic datasets

Two nuclear phylogenomic datasets were used for downstream analyses (Table S3). Dataset 1 comprised 3,023 orthologous loci, each present in at least 60 of the 97 sampled taxa. The concatenated nucleotide matrix contained 6,234,204 aligned sites. Of these, 2,370,130 sites (38.02%) were parsimony-informative. Missing characters accounted for 54.62% of the matrix. To assess the robustness of the nuclear phylogenetic backbone to character representation, the same 3,023 loci were also analyzed as amino acids (AA), first and second codon positions (NT12), and degeneracy-coded NT12 (NT12-deg). Dataset 2 comprised 678 orthologous loci with complete taxon occupancy across 10 representative taxa. Its concatenated nucleotide matrix contained 1,058,885 aligned sites, including 147,665 parsimony-informative sites (13.95%). Missing characters accounted for 22.37% of the matrix.

### 3.2 Phylogenetic results

The species tree inferred from Dataset 1 recovered all nine subfamilies of Dipsacales as monophyletic (Fig. 1). Adoxaceae was sister to Caprifoliaceae *s.l.* Within Caprifoliaceae *s.l.*, Diervilloideae diverged first, followed by Caprifolioideae. The remaining five subfamilies were resolved as (Morinoideae + Zabelioideae) sister to (Linnaeoideae + (Valerianoideae + Dipsacoideae)). All inter-subfamily backbone nodes received an LPP of 1.0. The same backbone topology was recovered in the concatenated ML analysis and in ASTRAL analyses based on AA, NT12, and NT12-deg data. All corresponding nodes received 100% bootstrap support in the concatenated ML tree and an LPP of 1.0 in the ASTRAL trees (Fig. S4–S7).

**Figure 1.**
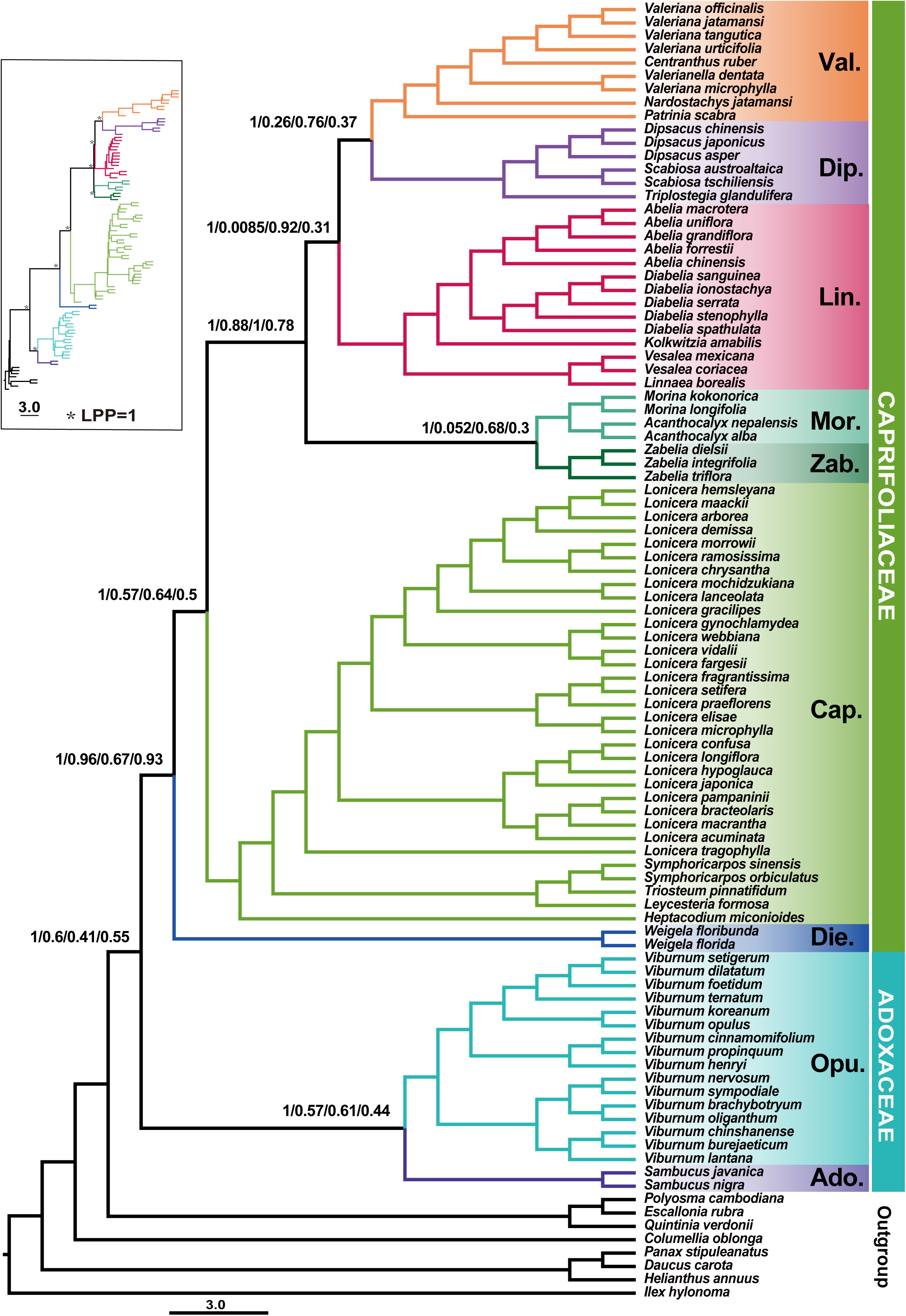
Nuclear ASTRAL species tree of Dipsacales inferred from Dataset 1 which consists of 3,023 orthologous nuclear loci. Values at major nodes indicate local posterior probability (LPP), quartet concordance (QC), quartet differential (QD) and quartet informativeness (QI), respectively and asterisks indicate LPP = 1. Branch and background colors identify subfamilies and families. The inset shows the complete tree. The scale bar represents coalescent units. Abbreviations: Val., Valerianoideae; Dip., Dipsacoideae; Lin., Linnaeoideae; Mor., Morinoideae; Zab., Zabelioideae; Cap., Caprifolioideae; Die., Diervilloideae; Opu., Opuloideae; Ado., Adoxoideae.

Despite these uniformly high support values, Quartet Sampling revealed substantial variation in concordance among backbone nodes (Fig. 1 and Fig. S8). The placement of Diervilloideae showed strong concordance (QC/QD/QI = 0.96/0.67/0.93), as did the five-subfamily clade following the divergence of Caprifolioideae (0.88/1.00/0.78). In contrast, concordance was weak for Morinoideae + Zabelioideae (0.052/0.68/0.30), (Linnaeoideae + (Valerianoideae + Dipsacoideae)) (0.0085/0.92/0.31), and Valerianoideae + Dipsacoideae (0.26/0.76/0.37).

The plastid ML tree also recovered all nine subfamilies as monophyletic, with 100% bootstrap support at all inter-subfamily nodes. However, the nuclear and plastid trees differed in the position of Linnaeoideae (Fig. 2). Nuclear analyses consistently recovered (Linnaeoideae + (Valerianoideae + Dipsacoideae)), whereas the plastid tree recovered Linnaeoideae + (Morinoideae + Zabelioideae).

**Figure 2.**
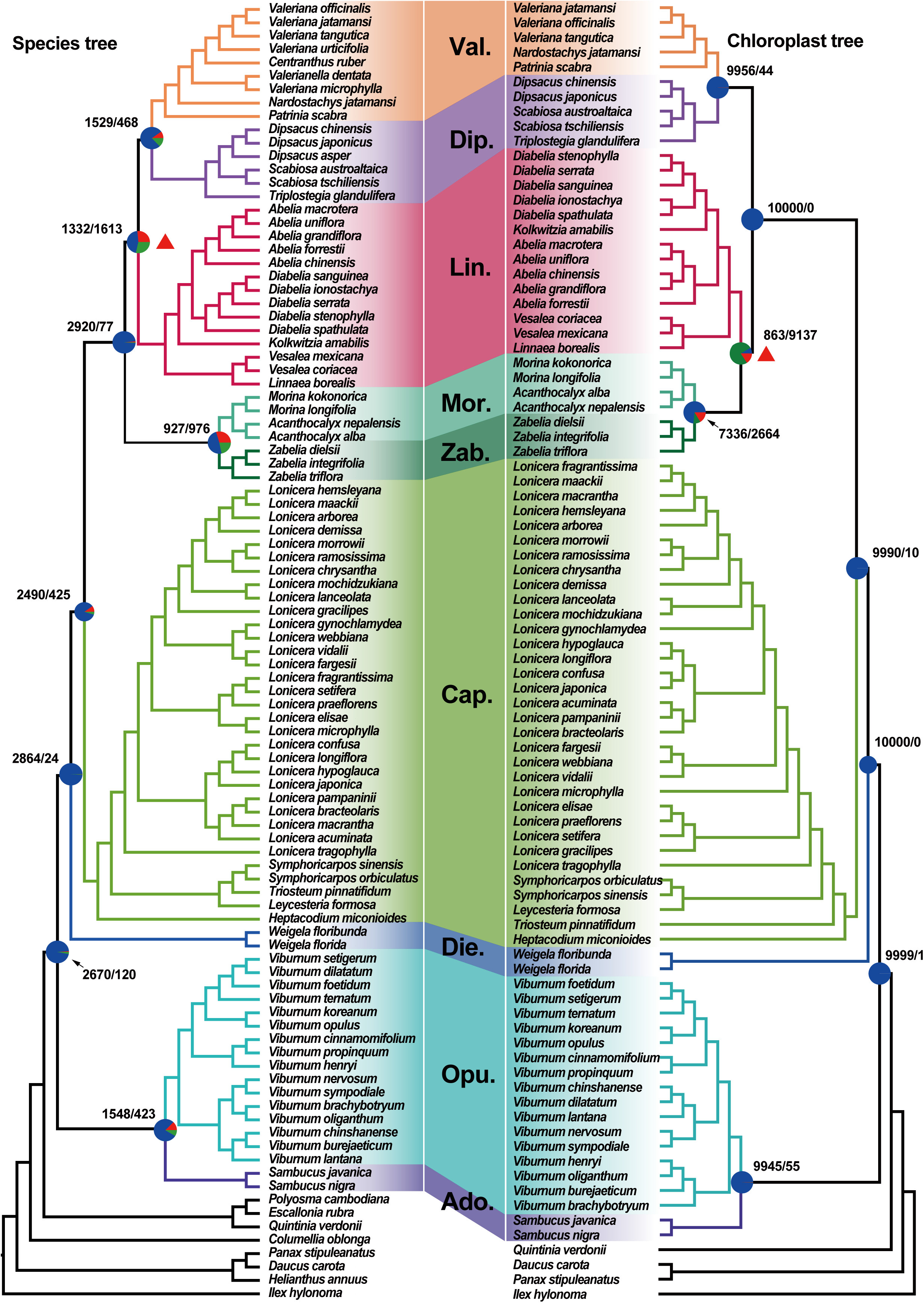
Phylogenetic conflict between the ASTRAL nuclear species tree on the left and the maximum-likelihood plastid tree of Dipsacales on the right. Colored bands connect corresponding taxa with pie charts on the nuclear tree show gene-tree support for the main and alternative topologies, and numbers indicate concordant/conflicting gene trees. Pie charts and numbers on the plastid tree summarize support/conflict in 10,000 ILS-only simulations. Red triangles mark the principal cytonuclear conflicts.

### 3.3 Gene-tree discordance and cytonuclear conflict

PhyParts revealed marked variation in gene-tree concordance across the inter-subfamily backbone (Fig. 2). The strongest conflict involved the placement of Linnaeoideae. At the node uniting Linnaeoideae with (Valerianoideae + Dipsacoideae), 1,332 gene trees supported the species-tree topology and 1,613 supported alternative topologies. (Morinoideae + Zabelioideae) also showed substantial conflict (927/976). (Valerianoideae + Dipsacoideae) showed greater concordance (1,529/468), and the deeper five-subfamily clade was strongly supported (2,920/77).

PhyTop revealed heterogeneous ILS and introgression/hybridization (IH) signals across backbone nodes (Fig. 3 and Fig. S9-S10). The strongest ILS signal occurred at (Linnaeoideae + (Valerianoideae + Dipsacoideae)) (N6; ILS index = 82.2%), followed by (Morinoideae + Zabelioideae) (N7; 65.1%). ILS accounted for 54.8% and 43.4% of the discordance at these two nodes, respectively. IH signals were generally weaker and highest at (Morinoideae + Zabelioideae) (N7; IH index = 22.6%; 7.9% explained).

**Figure. 3.**
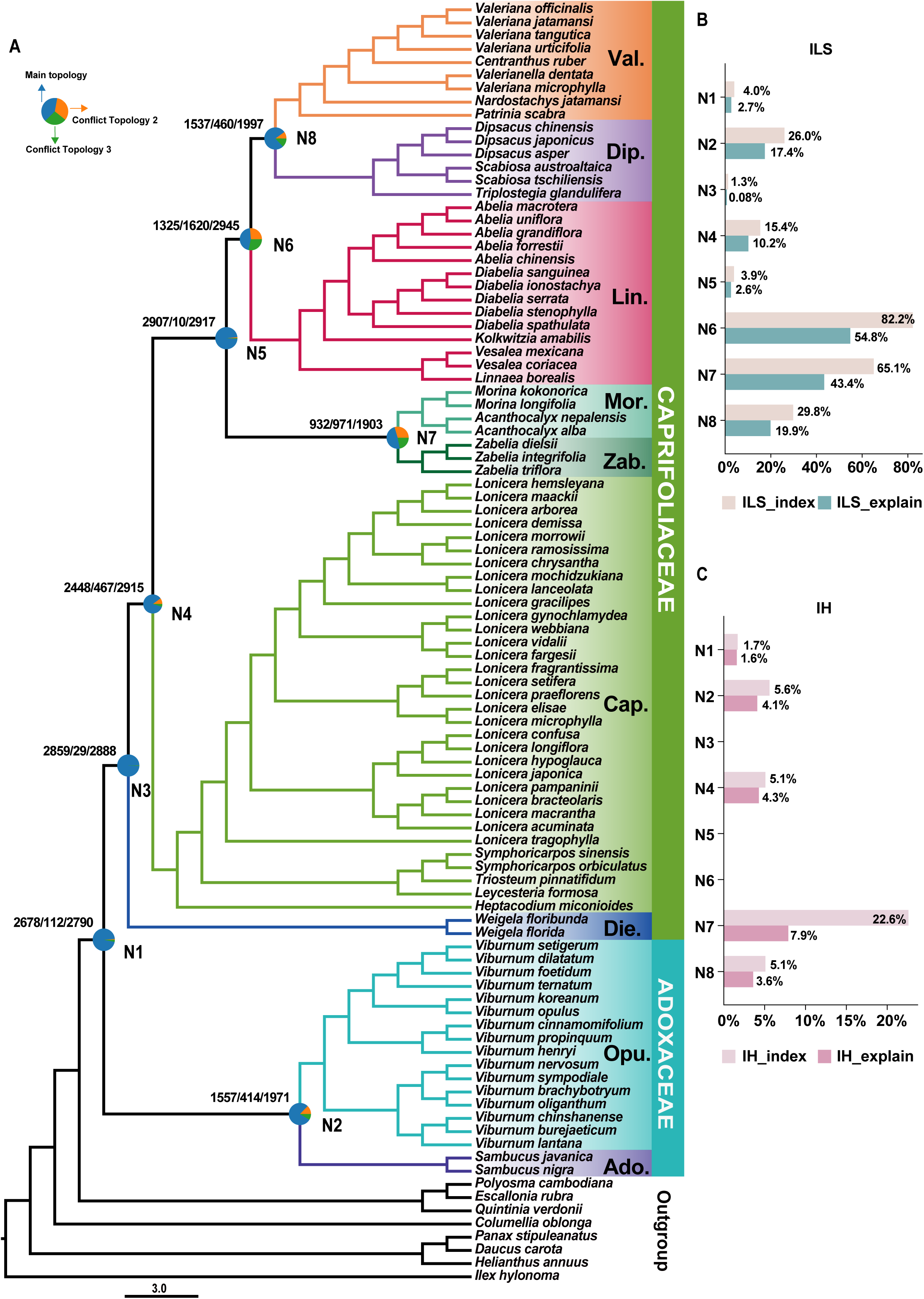
PhyTop estimates of ILS and introgression or hybridization associated with nuclear gene-tree conflict. A, ASTRAL species tree from Dataset 1. Pie charts show the main and two alternative topologies, and numbers indicate concordant/conflicting/informative gene trees. B, ILS_index and ILS_explain values. C, IH_index and IH_explain values. Nodes N1–N8 correspond to those in panel a.

Most plastid relationships were frequently recovered in the MSC simulations (Fig. 2). (Valerianoideae + Dipsacoideae) occurred in 9,956 of the 10,000 simulated trees, and (Morinoideae + Zabelioideae) in 7,336. Only 863 simulated trees recovered Linnaeoideae + (Morinoideae + Zabelioideae), while the remaining 9,137 supported alternative topologies.

### 3.4 Nuclear gene-tree discordance and MSC simulations

Dataset 2 recovered the same inter-subfamily backbone topology as Dataset 1 (Fig. 4A). For the node of (Linnaeoideae + (Valerianoideae + Dipsacoideae)) (N6), the main topology accounted for only 51.7% of gene trees, whereas the two alternative topologies accounted for 21.9% and 26.4%. A similar pattern was observed for (Morinoideae + Zabelioideae) (N7), with frequencies of 48.4%, 20.7%, and 30.9% for the three topologies, respectively. In contrast, the main topology predominated at several other backbone nodes. Its frequency reached 100% at N1, 99.1% at N3 and N5, and 85.9% at N4. The main topology was also more frequent at (Valerianoideae + Dipsacoideae) (N8; 78.4%) and (Opuloideae + Adoxoideae) (N2; 73.3%).

**Figure. 4.**
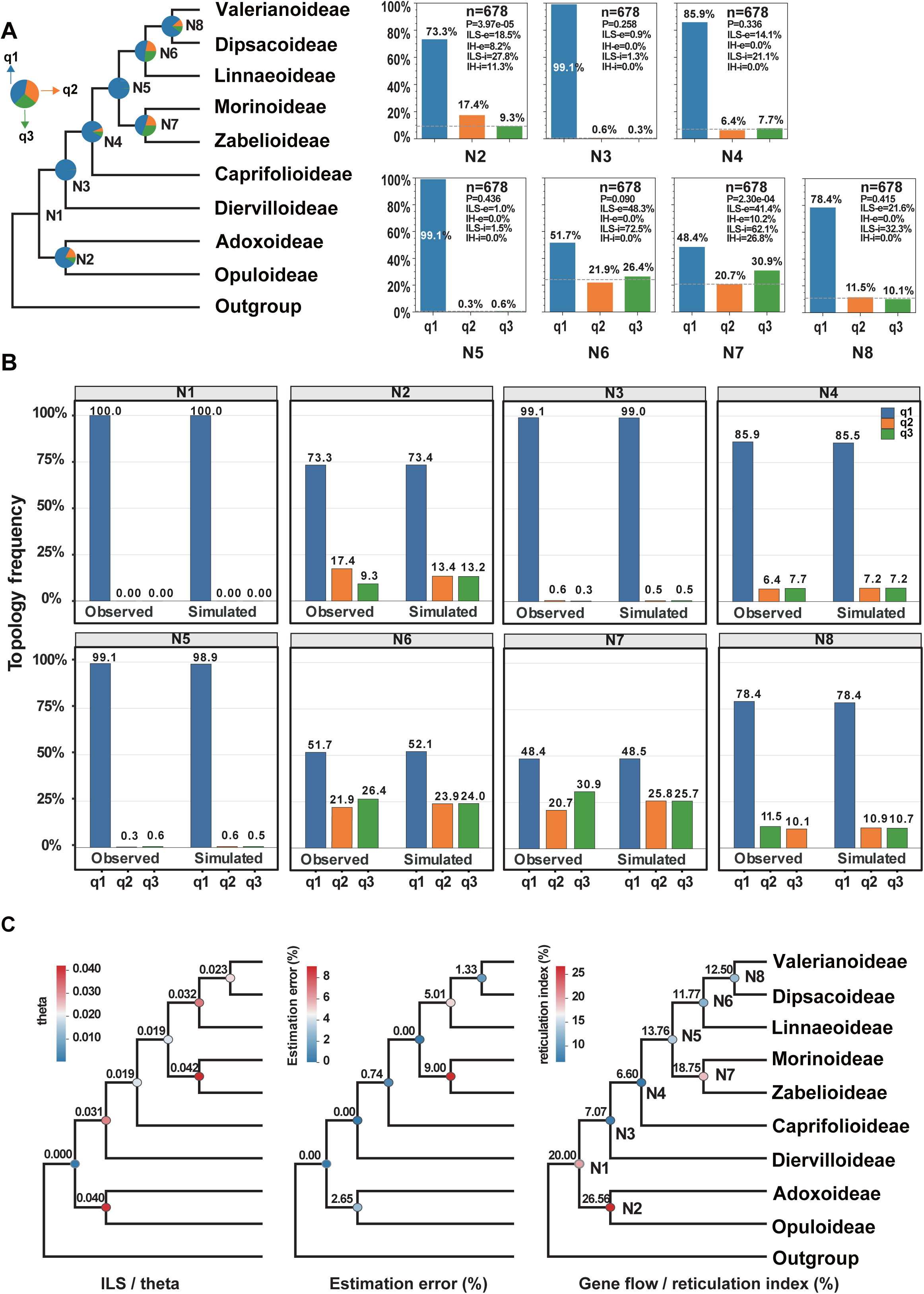
Potential sources of backbone discordance evaluated using Dataset 2. A, Ten-taxon species tree inferred from 678 completely occupied loci, with frequencies of topologies q1, q2, and q3. B, Comparison of observed topology frequencies with ILS-only coalescent simulations. C, Node-wise estimates of θ, gene-tree estimation error, and the reticulation index.

Under the ILS-only model, the two alternative topologies occurred at nearly equal frequencies in the MSC simulations (Fig. 4B). The empirical gene trees showed noticeable departures from this pattern at several nodes. For (Morinoideae + Zabelioideae) (N7), the two alternative topologies occurred at 20.7% and 30.9%, compared with 25.8% and 25.7% in the simulations. A similar asymmetry was observed for (Opuloideae + Adoxoideae) (N2; 17.4%/9.3% vs. 13.4%/13.2%). (Linnaeoideae + (Valerianoideae + Dipsacoideae)) (N6) showed a smaller deviation (21.9%/26.4% vs. 23.9%/24.0%).

### 3.5 Sources of nuclear gene-tree discordance

In Dataset 2, node-level estimates further revealed substantial variation in ILS, gene-tree estimation error, and reticulation signals (Fig. 4C). The highest θ value was detected for (Morinoideae + Zabelioideae) (N7; 0.042), followed by (Opuloideae + Adoxoideae) (N2; 0.040). (Linnaeoideae + (Valerianoideae + Dipsacoideae)) (N6) also showed a relatively high θ value of 0.032. Gene-tree estimation error was generally low but reached 9.00% at (Morinoideae + Zabelioideae) (N7) and 5.01% at (Linnaeoideae + (Valerianoideae + Dipsacoideae)) (N6). Reticulation indices ranged from 6.60% to 26.56%. The highest value occurred at (Opuloideae + Adoxoideae) (N2; 26.56%), followed by the Dipsacales crown node (N1; 20.00%) and (Morinoideae + Zabelioideae) (N7; 18.75%). The corresponding value for (Linnaeoideae + (Valerianoideae + Dipsacoideae)) (N6) was 11.77%.

QuIBL showed clear differences in model support among backbone nodes (Fig. 5). Among all discordant triplets, 86.3% were classified as ambiguous. The remaining 13.7% supported the ILS + Introgression model. No triplet supported the ILS-only model. The node-level patterns also differed. All triplets assigned to N2–N4 were ambiguous. At N1, 96.3% were ambiguous and 3.7% supported the ILS + Introgression model. At N5, 91.7% were ambiguous and 8.3% supported the ILS + Introgression model. All triplets assigned to N6 supported the ILS + Introgression model. At N7, 83.3% supported the ILS + Introgression model and 16.7% were ambiguous. At N8, the two categories were equally represented, with 50% supporting the ILS + Introgression model and 50% classified as ambiguous. Support for the ILS + Introgression model was therefore concentrated at N6 and N7. Most other backbone nodes were dominated by ambiguous classifications.

**Figure 5.**
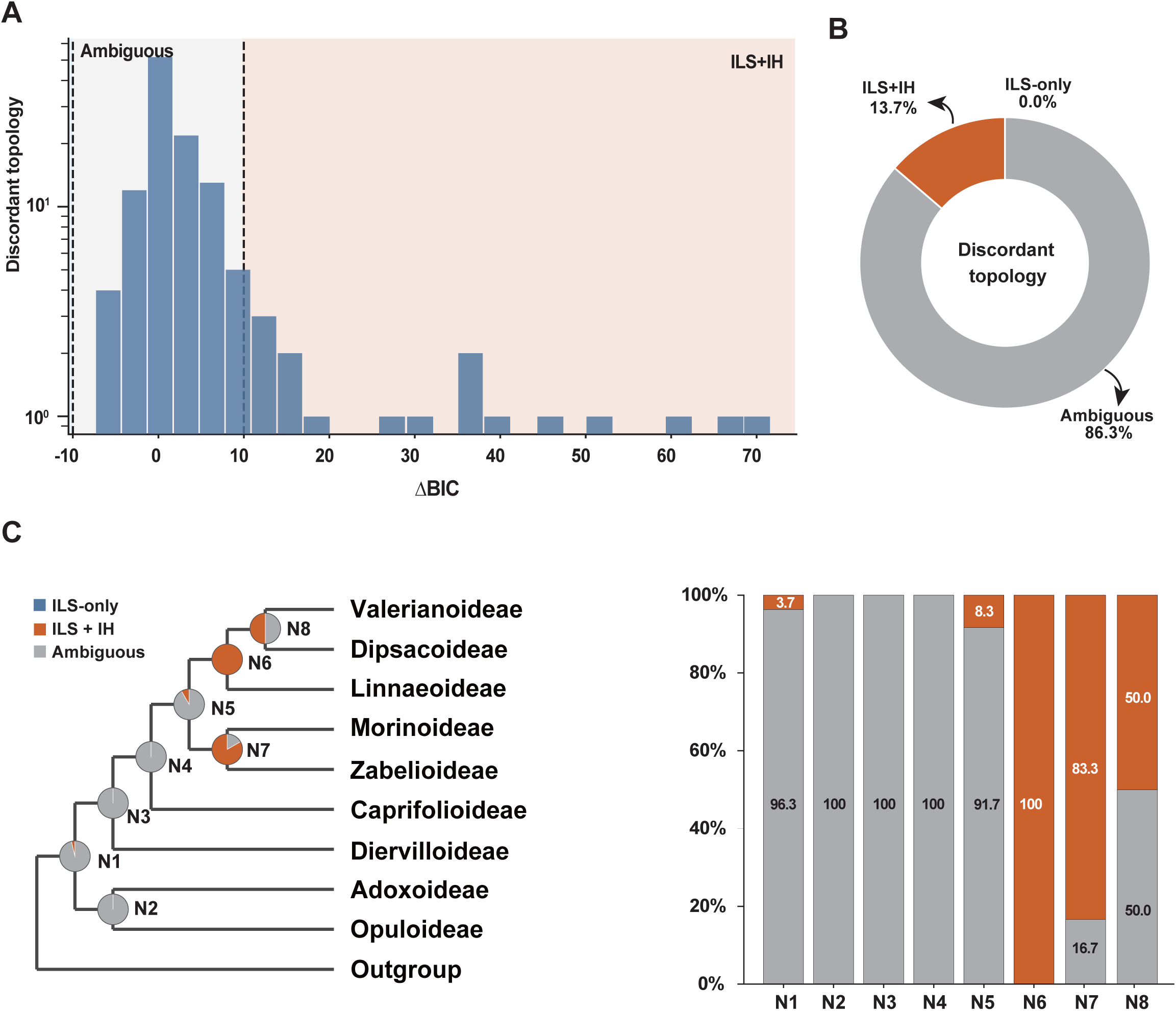
QuIBL model comparisons for discordant gene-tree topologies. (A) Distribution of ΔBIC values for discordant-topology records; the y-axis is logarithmic. ΔBIC = BIC_ILS-only_ − BIC_ILS + Introgression_. Dashed lines mark the classification thresholds: ΔBIC < −10 favors the ILS-only model, ΔBIC > 10 favors the ILS + Introgression model, and |ΔBIC| ≤ 10 is classified as ambiguous. (B) Overall proportions of records in the three model classes. (C) Proportions of records mapped to each species-tree node (N1–N8), shown as node pie charts and stacked bars.

### 3.6 Phylogenetic network inference

PhyloNet analyses showed that all network models with reticulations (H = 1-5) achieved higher the total log probability than the tree-only model (H = 0; Fig. 6B). The score increased in the one-reticulation model (H = 1; −12886.13) and reached maximum score in the two-reticulation model (H = 2; −12856.33). The three-, four-, and five-reticulation models had lower scores (H = 3; −12883.58), (H = 4; −12883.61), and (H = 5; −12872.59). Because allowing additional reticulations did not further improve the total log probability, the H = 2 network was selected as the best-scoring candidate network, which contained two inferred reticulation events (Fig. 6A). One involved Morinoideae; the two parental edges were associated with the lineage leading to (Linnaeoideae + (Valerianoideae + Dipsacoideae)) and the Zabelioideae lineage, with inheritance probabilities of 0.667 and 0.333. The second reticulation involved Adoxoideae and connected to two parental lineages: Opuloideae (γ = 0.915) and a deeper lineage within Caprifoliaceae *s.l.* (γ = 0.085).

**Figure 6.**
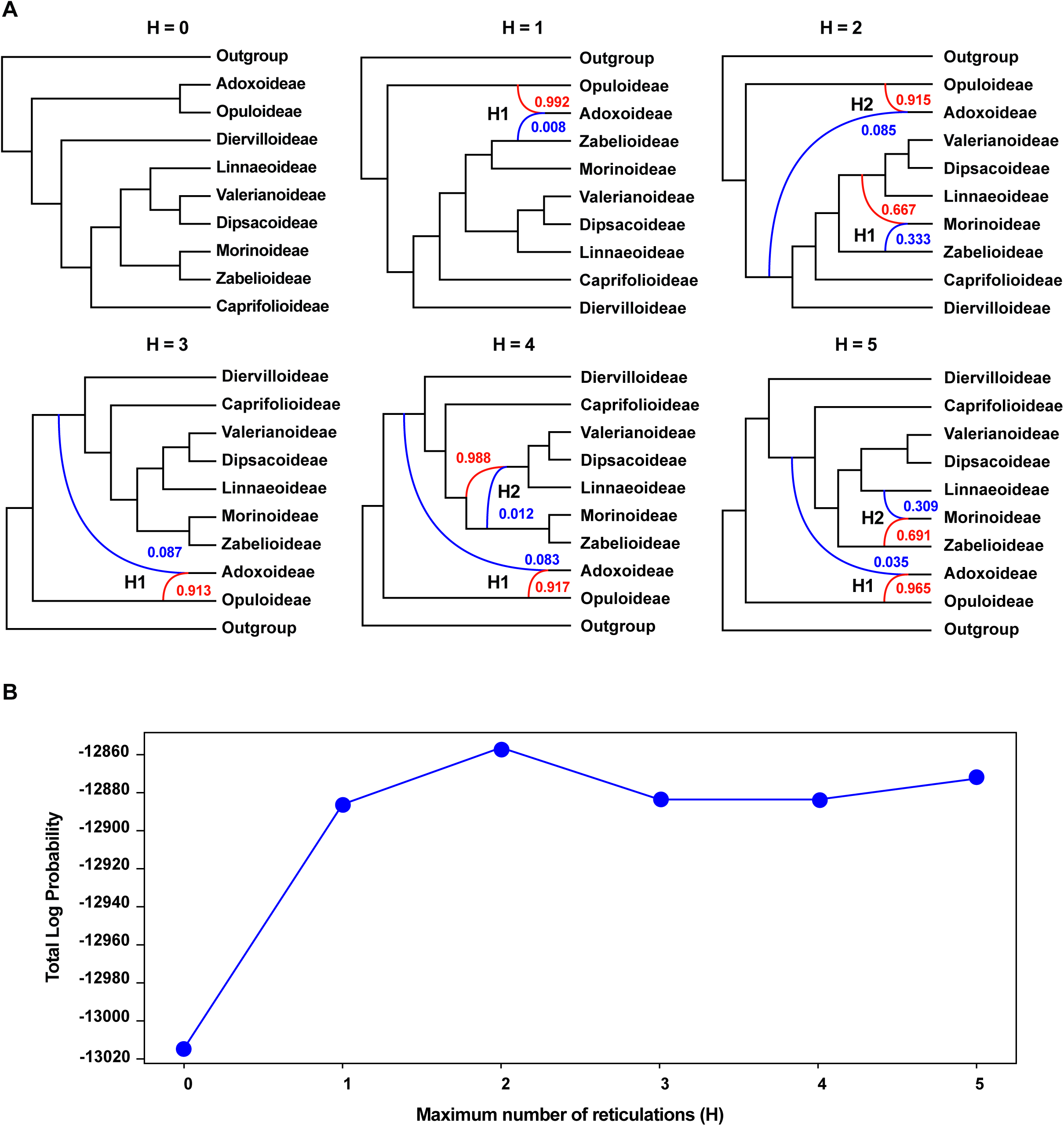
PhyloNet inference of reticulate evolution among Dipsacales subfamilies. A, Maximum-pseudolikelihood networks allowing zero to five reticulations. Colored reticulation edges indicate parental contributions, with inheritance probabilities shown beside the edges. B, Best total log probability across 200 searches for H = 0–5. H = 2 network was retained as the least-complex candidate beyond which additional reticulations did not improve the score.

### 3.7 Divergence time estimation

Divergence-time analysis of the nuclear dataset placed the crown of Dipsacales in the Late Cretaceous at 83.63 Ma (95% HPD = 71.94–95.86 Ma; Fig. 7). The crown of Caprifoliaceae, defined by the split of Diervilloideae from the remaining six subfamilies, was dated to 70.61 Ma (95% HPD = 59.69–82.15 Ma). Within the resulting six-subfamily clade, Caprifolioideae diverged from the other five subfamilies at 65.21 Ma (95% HPD = 53.92–75.89 Ma). The split between Opuloideae and Adoxoideae was dated to 59.07 Ma (95% HPD = 48.11–74.97 Ma).

**Figure 7.**
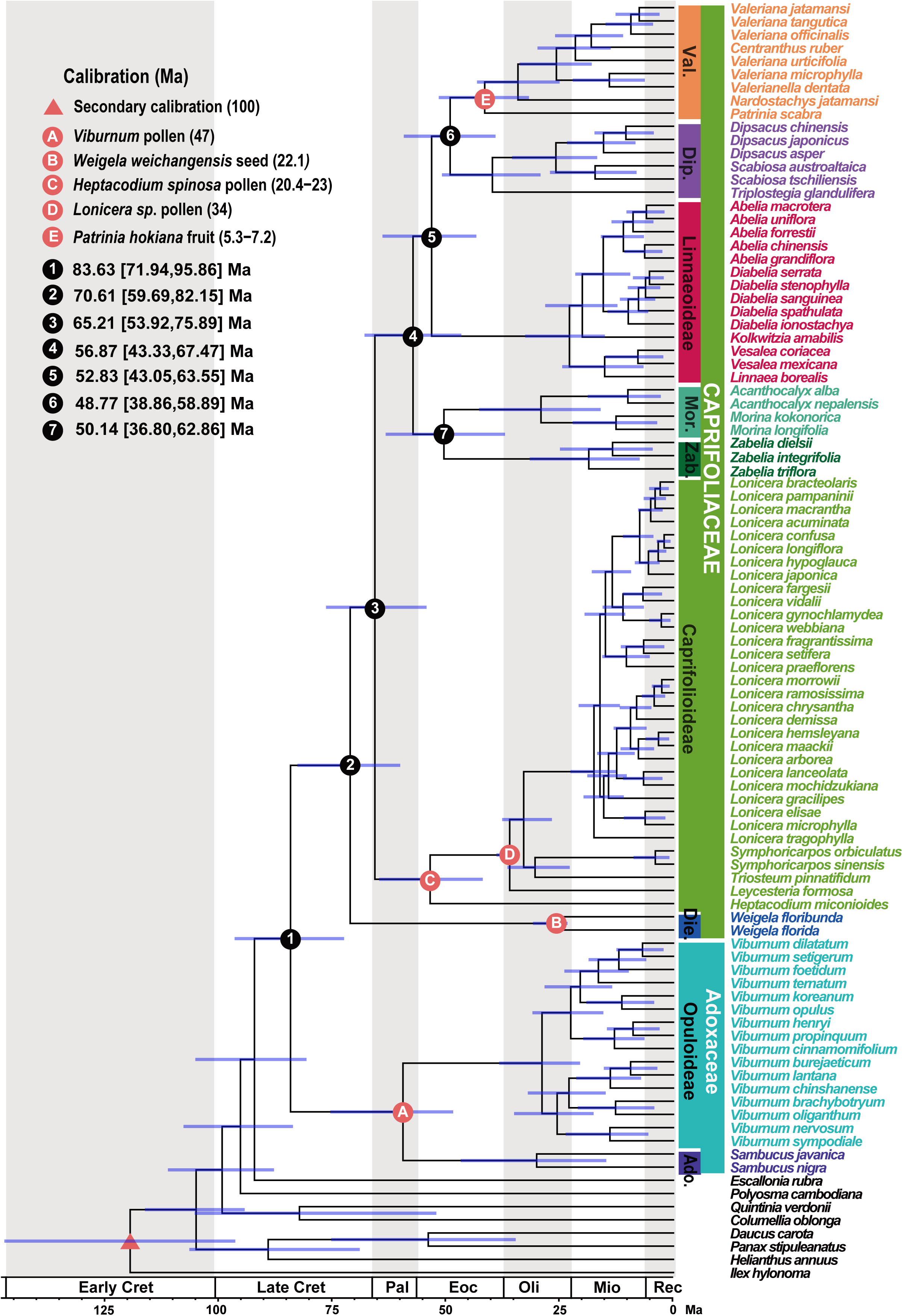
Divergence-time estimation of Dipsacales based on nuclear data. The chronogram represents the maximum clade credibility tree, with blue bars indicating 95% highest posterior density intervals. The red triangle marks the secondary calibration, letters A–F indicate fossil calibrations, and numbered black circles identify major divergence events and their estimated ages. Colored bars identify subfamilies and families. Ma, million years ago.

Within the five-subfamily clade, point estimates for the backbone divergences fell in the late Paleocene and early Eocene. (Morinoideae + Zabelioideae) diverged from (Linnaeoideae + (Valerianoideae + Dipsacoideae)) at 56.87 Ma (95% HPD = 43.33–67.47 Ma). Linnaeoideae split from (Valerianoideae + Dipsacoideae) at 52.83 Ma (95% HPD = 43.05–63.55 Ma). Morinoideae and Zabelioideae diverged at 50.14 Ma (95% HPD = 36.80–62.86 Ma), and Valerianoideae and Dipsacoideae diverged at 48.77 Ma (95% HPD = 38.86–58.89 Ma).

Crown ages were estimated for all nine subfamilies of Dipsacales. Caprifolioideae was dated to 53.15 Ma (95% HPD = 41.65–64.28 Ma). Valerianoideae and Dipsacoideae had crown ages of 41.23 Ma (95% HPD = 31.56–51.27 Ma) and 39.55 Ma (95% HPD = 29.04–50.57 Ma), respectively. The crown age of Adoxoideae was 29.87 Ma (95% HPD = 14.67–46.47 Ma). Morinoideae and Opuloideae had crown ages of 28.94 Ma (95% HPD = 15.90–42.45 Ma) and 28.70 Ma (95% HPD = 20.39–38.07 Ma), respectively. Diervilloideae was dated to 25.91 Ma (95% HPD = 23.13–30.67 Ma). Linnaeoideae and Zabelioideae had crown ages of 22.71 Ma (95% HPD = 14.97–32.38 Ma) and 18.48 Ma (95% HPD = 7.37–31.39 Ma), respectively.

## 4 Discussion

### 4.1 A stable nuclear backbone despite widespread gene-tree discordance

Our analyses included 89 species representing all nine Dipsacales subfamilies and 3,023 orthologous nuclear loci. They recovered (Morinoideae + Zabelioideae) as sister to (Linnaeoideae + (Valerianoideae + Dipsacoideae)). This backbone was consistently recovered by concatenation and ASTRAL analyses (Fig. 1 and Fig. S4). It also remained unchanged across amino acid, NT12 and degeneracy-recoded NT12 character representations (Fig. S5–S7). This topology agrees with the concatenation analysis of Wang et al. (2021) and the exon-only ASTRAL analysis of Lee et al. (2021). It also matches the Dipsacales subtree in the angiosperm species tree of Zuntini et al. (2024). These results further support Zabelia as sister to Morinoideae and Linnaeoideae as sister to the (Valerianoideae + Dipsacoideae) clade.

Previous differences in the placement of *Zabelia* and the monophyly of Dipsacoideae may reflect data composition and how inference methods summarize conflicting signals. In Lee et al. (2021), adding flanking regions increased both alignment length and the proportion of informative sites. However, the position of *Zabelia* also changed in the ASTRAL analysis. Thus, adding characters may alter relative support for alternative topologies as well as increase phylogenetic information (Shen et al., 2017). Concatenation estimates a topology from the combined sequence alignment across loci. ASTRAL instead infers a species tree from quartet relationships in the input gene trees (Mirarab et al., 2014). Under extensive genealogical heterogeneity, differences in locus informativeness and gene-tree estimation accuracy may contribute to different relationships recovered by these methods (Roch and Warnow, 2015). Differences among previous studies may therefore reflect variation in nuclear-locus sampling and informativeness, as well as how inference methods integrate conflicting genealogical signals. Across 3,023 nuclear loci, our concatenation, ASTRAL and character-recoding analyses consistently recovered the same relationships among the five subfamilies. This consistency strengthens support for the topology as a representation of the major lineage divergences within Linnina.

These results highlight the value of using more nuclear loci to resolve deep phylogenetic relationships in groups with extensive gene-tree discordance. Including more high-quality loci may better capture variation among gene genealogies and reduce uncertainty associated with analyses based on few loci. These data can help clarify relationships among major lineages and assess the evolutionary processes underlying gene-tree discordance. Therefore, developing taxon-specific nuclear-locus reference sets to expand the range of orthologous loci analyzed is a strategy that merits wider use in phylogenomic studies of complex groups.

### 4.2 Processes underlying deep phylogenetic discordance in Dipsacales

To investigate deep gene-tree discordance in Linnina, we constructed Dataset 2 by optimizing representative-taxon selection for subfamily coverage and sequence completeness. PhyTop analyses showed similar patterns of incomplete lineage sorting (ILS) and introgression/hybridization (IH) at N6 (Linnaeoideae + (Valerianoideae + Dipsacoideae)) and N7 (Morinoideae + Zabelioideae) in both datasets (Figs. 3–4). Despite the limited taxon sampling in Dataset 2, this consistency provides support for our qualitative interpretation of discordance at these nodes. Our results suggest that ILS underlies much of the deep nuclear gene-tree discordance in Linnina. Signals of ancient gene flow nevertheless differ among the ancestral branches examined. Parametric simulations following Cai et al. (2021) indicated that gene-tree estimation error alone cannot explain the major patterns of gene-tree discordance at nodes N6 (Linnaeoideae + (Valerianoideae + Dipsacoideae)) and N7 (Morinoideae + Zabelioideae) (Fig. 4C). At N6 (Linnaeoideae + (Valerianoideae + Dipsacoideae)), topology frequencies approached ILS-only expectations, whereas internal branch-length distributions supported a model incorporating gene flow. At N7 (Morinoideae + Zabelioideae), evidence for gene flow was more consistent across topology frequencies, branch-length distributions and network inference (Fig. 4–6). This contrast suggests that ancient gene flow can affect gene coalescence histories without producing significant asymmetry between the two minor topology frequencies. In summary, our results indicate that ILS is the predominant source of deep nuclear gene-tree discordance in Linnina, while additional evidence for ancient gene flow varies among nodes. The detection of ancient gene flow depends partly on how its signals are reflected in topology frequencies and internal branch-length distributions.

The different signals at N6 (Linnaeoideae + (Valerianoideae + Dipsacoideae)) and N7 (Morinoideae + Zabelioideae) may help explain contrasting assessments of ancient gene flow in previous studies. Wang et al. (2021) inferred candidate reticulation events within Linnina, whereas the SNAQ analysis of Lee et al. (2021) detected no clear introgression signal. These differences may reflect variation in taxon sampling, locus informativeness, and the statistical features used to detect ancient gene flow. Topology frequencies and internal branch lengths capture different aspects of gene genealogies (Edelman et al., 2019). Strong ILS can reduce frequency differences between the two minor topologies, thereby reducing the ability of topology-asymmetry tests to detect gene flow (Shang et al., 2025). Limited information in conserved loci may also hinder the detection of ancient events (Lee et al., 2021). In our analyses, topology frequencies and branch lengths provided complementary evidence at N6 (Linnaeoideae + (Valerianoideae + Dipsacoideae)), whereas the signals at N7 (Morinoideae + Zabelioideae) were more consistent across methods. We therefore suggest that future studies integrate topology frequencies and internal branch-length distributions when assessing potential ancient gene flow. Considering both sources of evidence may improve the detection and interpretation of ancient gene flow signals, particularly in the presence of strong ILS.

Nuclear data place Linnaeoideae as sister to Valerianoideae + Dipsacoideae, whereas plastid data support its sister relationship with Morinoideae + Zabelioideae, consistent with previous reports of cytonuclear discordance (Wang et al., 2021; Lee et al., 2021). In our ILS-only simulations, with coalescent branch lengths adjusted to reflect the assumed plastid inheritance model, 8.63% of simulated trees recovered the observed plastid topology (Fig. 2). This result indicates that the plastid placement can arise under ILS alone. However, nuclear evidence of ancient gene flow involving the relevant ancestral branches also suggests plastid capture as a possible explanation. Introgression has been implicated in cytonuclear discordance in annual sunflowers and *Dipelta* (Lee-Yaw et al., 2019; Cao et al., 2025). Although the observed plastid placement of Linnaeoideae can arise under ILS alone, its relatively low frequency in the simulations, together with the inferred ancestral reticulation, raises the possibility that ancient gene flow, potentially involving plastid capture, contributed to the cytonuclear discordance.

Divergence-time estimates and historical biogeography provide temporal and geographical context for the discordance at N6 (Linnaeoideae + (Valerianoideae + Dipsacoideae)) and N7 (Morinoideae + Zabelioideae). The two major Linnina clades diverged approximately 56.87 Ma (Fig. 7). Linnaeoideae subsequently diverged from Valerianoideae + Dipsacoideae at approximately 52.83 Ma, whereas Morinoideae diverged from Zabelioideae at approximately 50.14 Ma. The corresponding differences between node-age point estimates are 4.04 and 6.73 Myr, respectively. These successive divergences provide a temporal context for the persistence of ancestral polymorphism. If the intervals were short relative to ancestral coalescent times, ancestral polymorphism could have persisted across successive speciation events (Pamilo and Nei, 1988). This scenario is consistent with the substantial ILS signals at both nodes. Evidence of ancient gene flow further suggests that the separation of some ancestral lineages may have been accompanied by genetic exchange. Previous biogeographical studies linked the early evolution of Morinoideae, Valerianoideae and Dipsacoideae to Asia and highlighted East Asia in early Linnaeoideae evolution. These findings provide geographical indications of potential contact among some ancestral lineages (Bell and Donoghue, 2003; Wang et al., 2015). Together with our results, this interpretation suggests that early Linnina divergence may have involved two overlapping processes. Successive lineage divergences may have allowed ancestral polymorphism to persist, while continued or renewed gene flow further modified some gene genealogies. This scenario could explain the shared ILS background at N6 (Linnaeoideae + (Valerianoideae + Dipsacoideae)) and N7 (Morinoideae + Zabelioideae) alongside their different gene-flow signals. Further work should determine whether gene flow occurred during lineage separation or through later secondary contact, and identify the ancestral lineages and regions involved.

### 4.3 Implications of genealogical heterogeneity for morphological evolution in Dipsacales

Our analyses consistently recovered Morinoideae + Zabelioideae as one major Linnina clade and Linnaeoideae + (Valerianoideae + Dipsacoideae) as the other. This backbone further supports previous interpretations of shared pollen characters and the evolution of the herbaceous habit. The psilate exine and continuous ring-shaped endoaperture shared by Morinoideae and Zabelioideae may have been inherited from their common ancestor (Jacobs et al., 2011). By contrast, the herbaceous habit may have evolved separately in Morinoideae and the Valerianoideae + Dipsacoideae clade (Lee et al., 2021). Our analyses of deep discordance raise a further possibility: separate origins of a phenotype need not imply independent origins of all contributing genetic variants (Lee and Coop, 2017). The substantial ILS background and evidence of ancient gene flow at N6 (Linnaeoideae + (Valerianoideae + Dipsacoideae)) and N7 (Morinoideae + Zabelioideae) provide a historical context for examining this possibility.

Stamen reduction provides a concrete example for examining this distinction. Our nuclear backbone, considered together with character distributions among related taxa, favors separate reductions to two fertile stamens in *Morina* and *Fedia*. *Acanthocalyx* retains four stamens, whereas *Morina* has two fertile stamens and a second pair of staminodes. Three stamens are common in core Valerianoideae, and the two-stamen condition in *Fedia* has been interpreted as a further reduction (Donoghue et al., 2003). Both *Morina* and *Fedia* retain the dorsal pair of fertile stamens. However, two stamen primordia were observed during early floral development in *Fedia graciliflora* (Berger et al., 2021). Retention of fertile stamens in corresponding positions may therefore involve changes at different developmental stages. These developmental differences do not, by themselves, establish the historical origins of the underlying functional variants. Wang et al. (2021) proposed that differences in stamen number within Morinoideae might involve introgressed alleles. The ancestral histories inferred here motivate a further, testable hypothesis. Stamen reduction may have occurred separately in *Morina* and *Fedia*, while some contributing functional variants may predate the corresponding phenotypic transitions. These variants could have persisted from ancestral populations or spread through ancient gene flow among ancestral lineages. They could subsequently have contributed to changes in stamen development under lineage-specific regulatory contexts.

The contribution of older genetic variation to later morphological specialization has been demonstrated in other plants. In *Capsella rubella*, reduced petal size partly involves regulatory variation in a petal-specific enhancer of *SAP* (Sicard et al., 2016). In the *Mimulus aurantiacus* species complex, an ancient introgressed haplotype containing *MaMyb2* persisted in descendant lineages and contributed to the repeated evolution of red flowers (Stankowski and Streisfeld, 2015; Short and Streisfeld, 2023). These examples show that phenotypic transitions can postdate the origin of their contributing functional variants. Testing this hypothesis in Linnina would require comparing stamen development in *Morina*, *Fedia*, and their relatives with higher numbers of fertile stamens. Identifying candidate genetic variants, examining their functional effects, and reconstructing local gene genealogies could help determine whether similar reductions in stamen number involve independent genetic changes or shared ancestral variation. If shared variants are identified, broader genomic sampling and further analyses would be needed to distinguish their retention through ILS from their transfer through ancient introgression.

Candidate reticulation signals at node N2 (Adoxoideae + Opuloideae) provide additional insights into the morphological and ecological divergence of *Viburnum* and *Sambucus*. Lens et al. (2016) linked differences in vessel perforation plates between *Viburnum* (scalariform) and *Sambucus* (simple) to ancestral habitats and hydraulic function. Landis et al. (2021) further suggested that *Viburnum* initially diversified in warm-temperate forests of Asia. The candidate reticulation signals detected in our study suggest that ancient gene flow may have occurred among the relevant ancestral lineages. Their morphological and ecological divergence may therefore not have occurred under complete genetic isolation. However, direct evidence linking ancient gene flow to the evolution of these traits is still lacking. Future studies should integrate genealogies of relevant functional loci, the temporal and geographic distributions of ancestral lineages, and analyses of character evolution to test whether retained ancestral variation or ancient gene flow contributed to morphological divergence in these lineages.

## 5 Conclusions

Our analyses of 3,023 orthologous nuclear loci consistently recovered the same phylogenetic backbone of Dipsacales across inference methods and character-coding schemes, despite extensive gene-tree discordance. Node-level analyses suggest that incomplete lineage sorting (ILS) is a major source of deep discordance in Linnina, whereas evidence for ancient gene flow differs among ancestral branches. The plastid placement of Linnaeoideae could arise under ILS alone, although ancient introgression, potentially involving plastid capture, remains a possible additional explanation. Divergence-time estimates indicate that the major Linnina lineages diverged in relatively close succession during the late Paleocene and early Eocene. Overall, our study further clarifies relationships among the major lineages of Dipsacales. It also improves our understanding of how ILS and ancient gene flow jointly contribute to deep phylogenetic discordance during rapid successive divergences.

**Figure S1.**
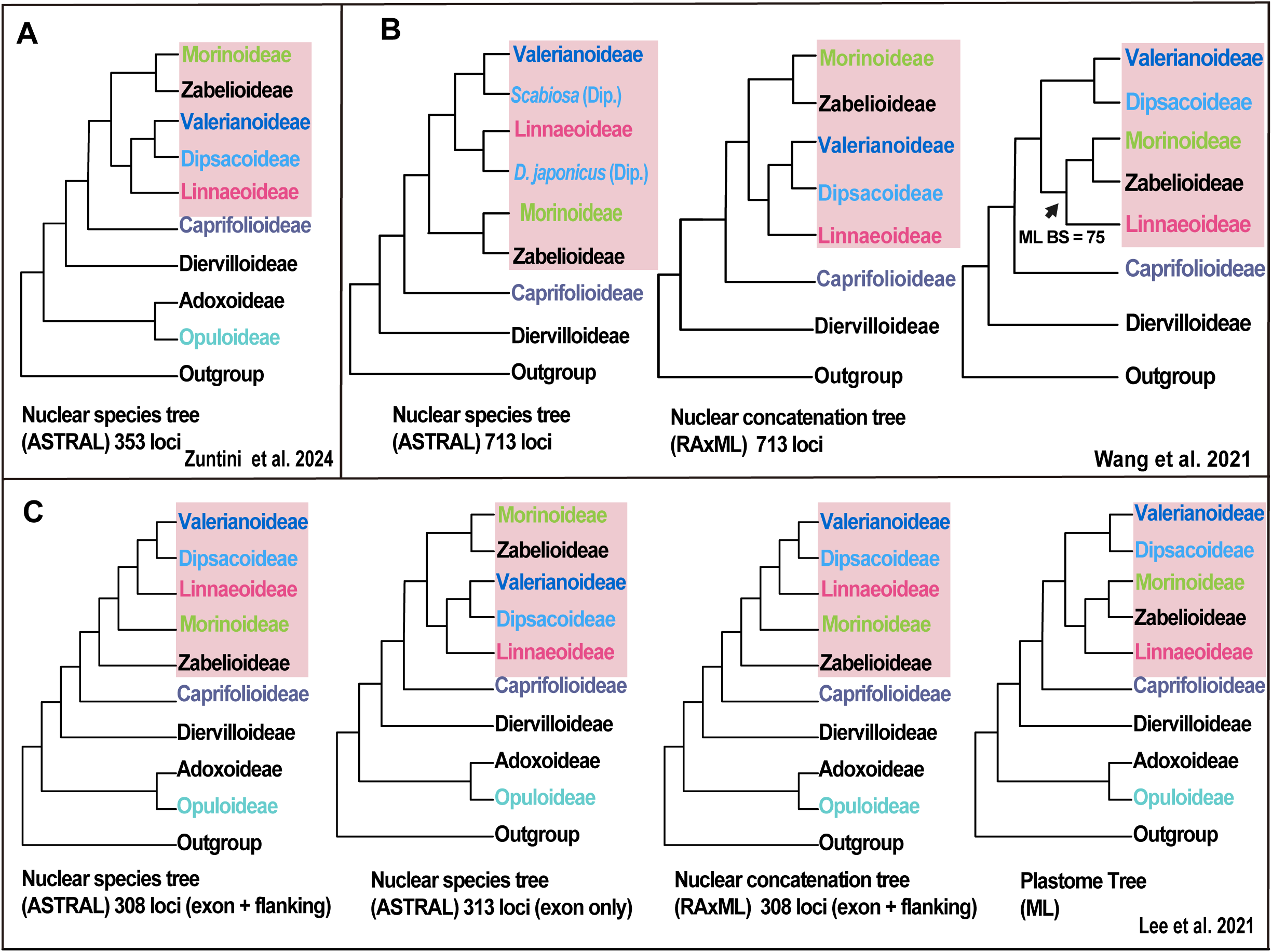
Phylogenetic relationships among Dipsacales subfamilies inferred from different genomic datasets and analytical approaches in previous studies. Topologies were simplified from the original studies to illustrate subfamily-level relationships.

**Figure S2.**
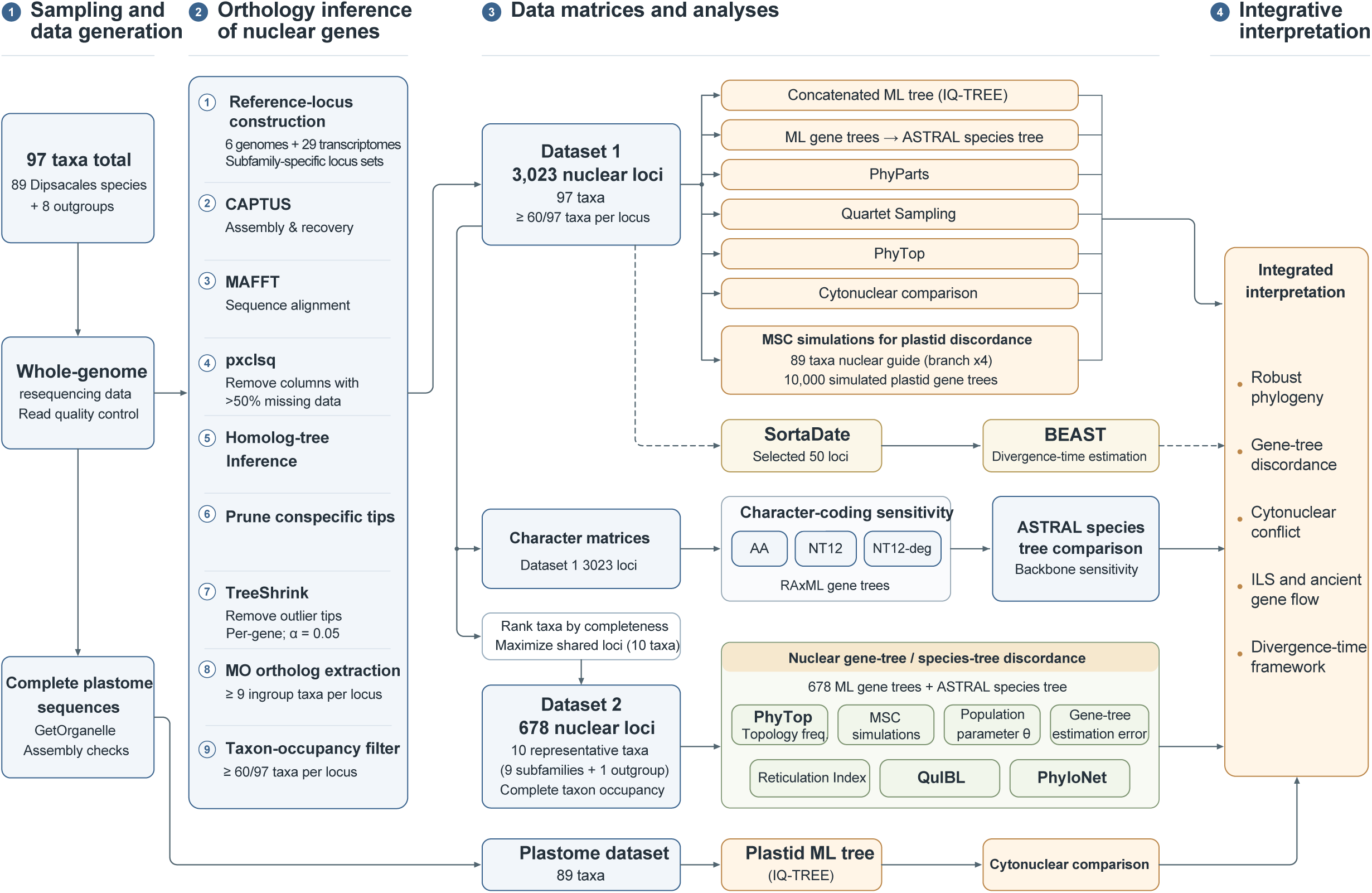
Overview of the analytical workflow used in this study. The diagram summarizes taxon sampling, nuclear locus recovery and orthology inference, construction of the two nuclear datasets, and their applications in phylogenetic inference, gene-tree discordance analyses, phylogenetic network inference, and divergence-time estimation.

**Figure S3.**
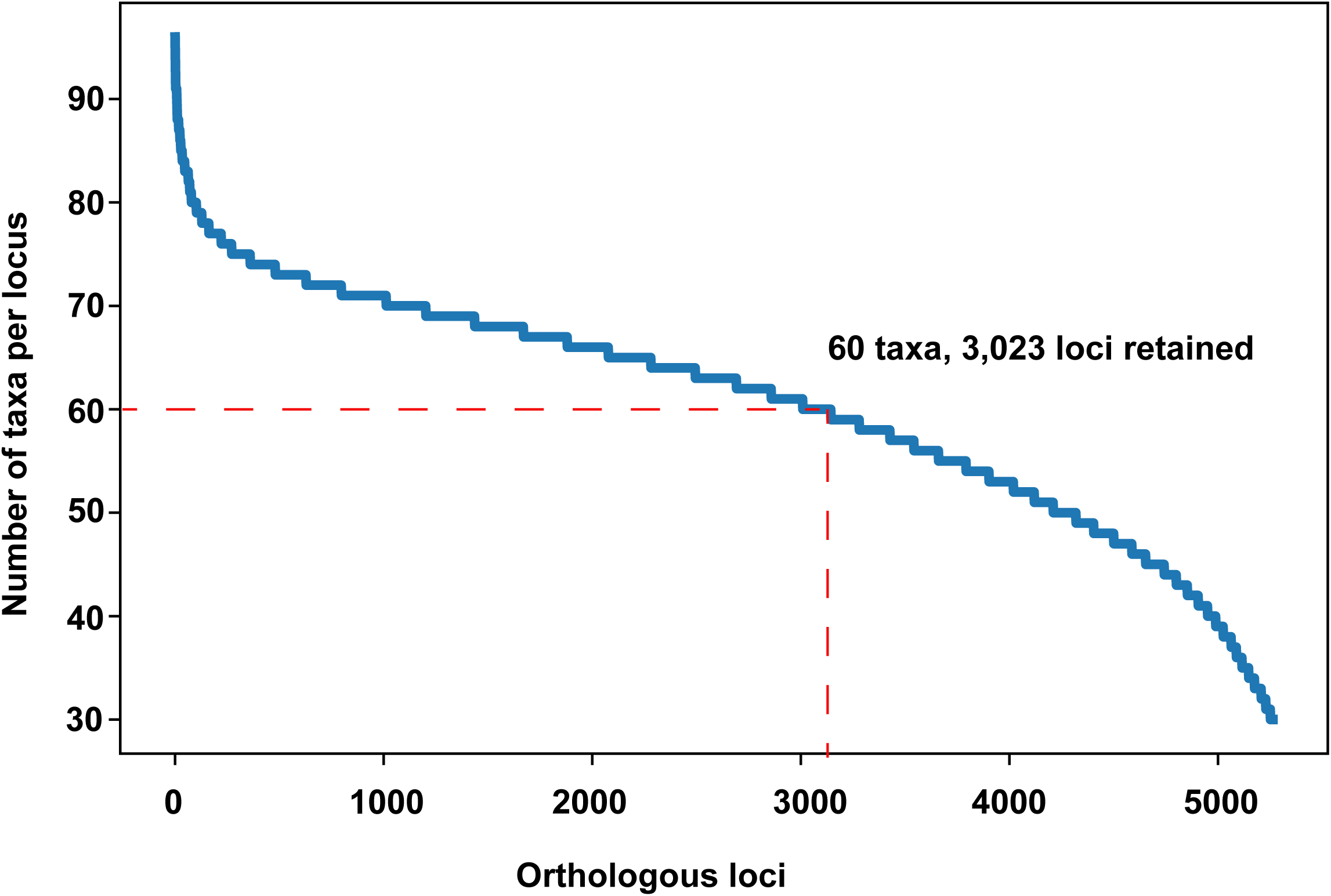
Taxon occupancy distribution of orthologous nuclear loci. The x-axis shows the ranked orthologous loci, and the y-axis shows the number of taxa represented per locus. The red dashed lines indicate the occupancy threshold of 60 taxa.

**Figure S4.**
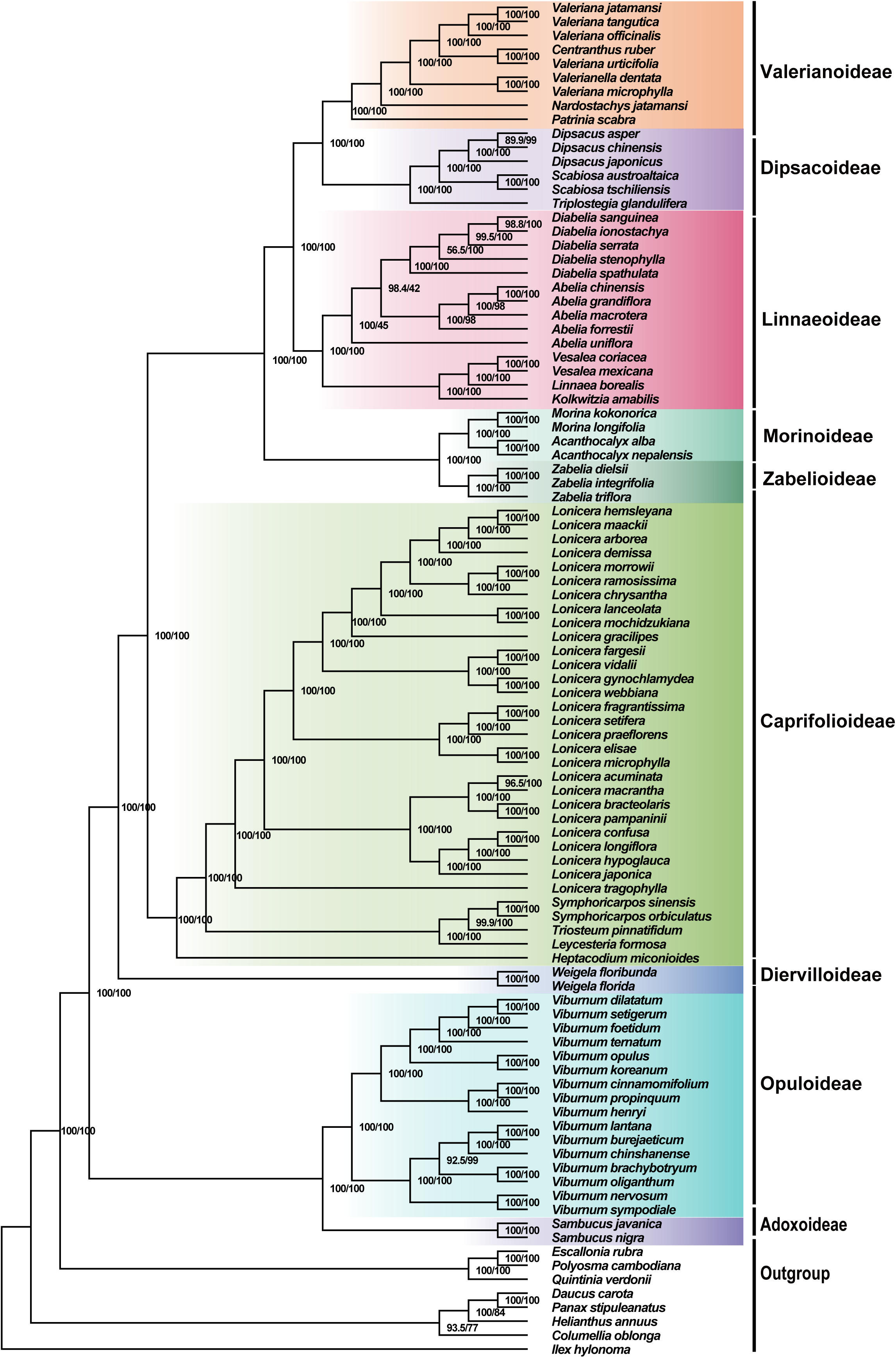
Maximum-likelihood tree of Dipsacales inferred from the partitioned concatenated matrix of Dataset 1. The maximum-likelihood tree was inferred in IQ-TREE from 3,023 orthologous nuclear loci. Branch support values are shown as SH-aLRT/UFBoot. Background colors and labels identify the nine subfamilies.

**Figure S5.**
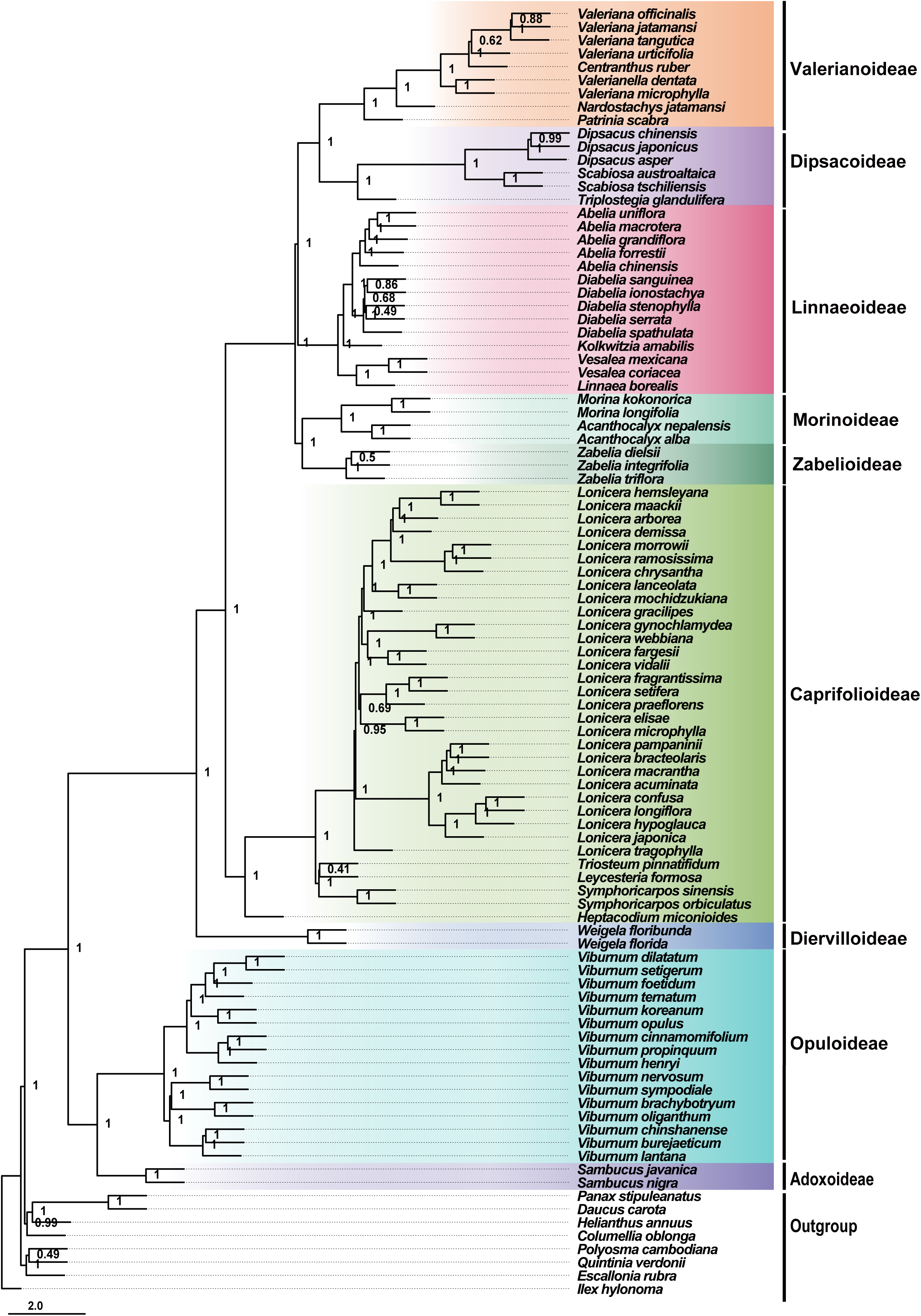
ASTRAL species tree of Dipsacales based on amino-acid data derived from Dataset 1. Node labels show local posterior probabilities (LPPs). Background shading denotes subfamilies. Branch lengths are measured in coalescent units.

**Figure S6.**
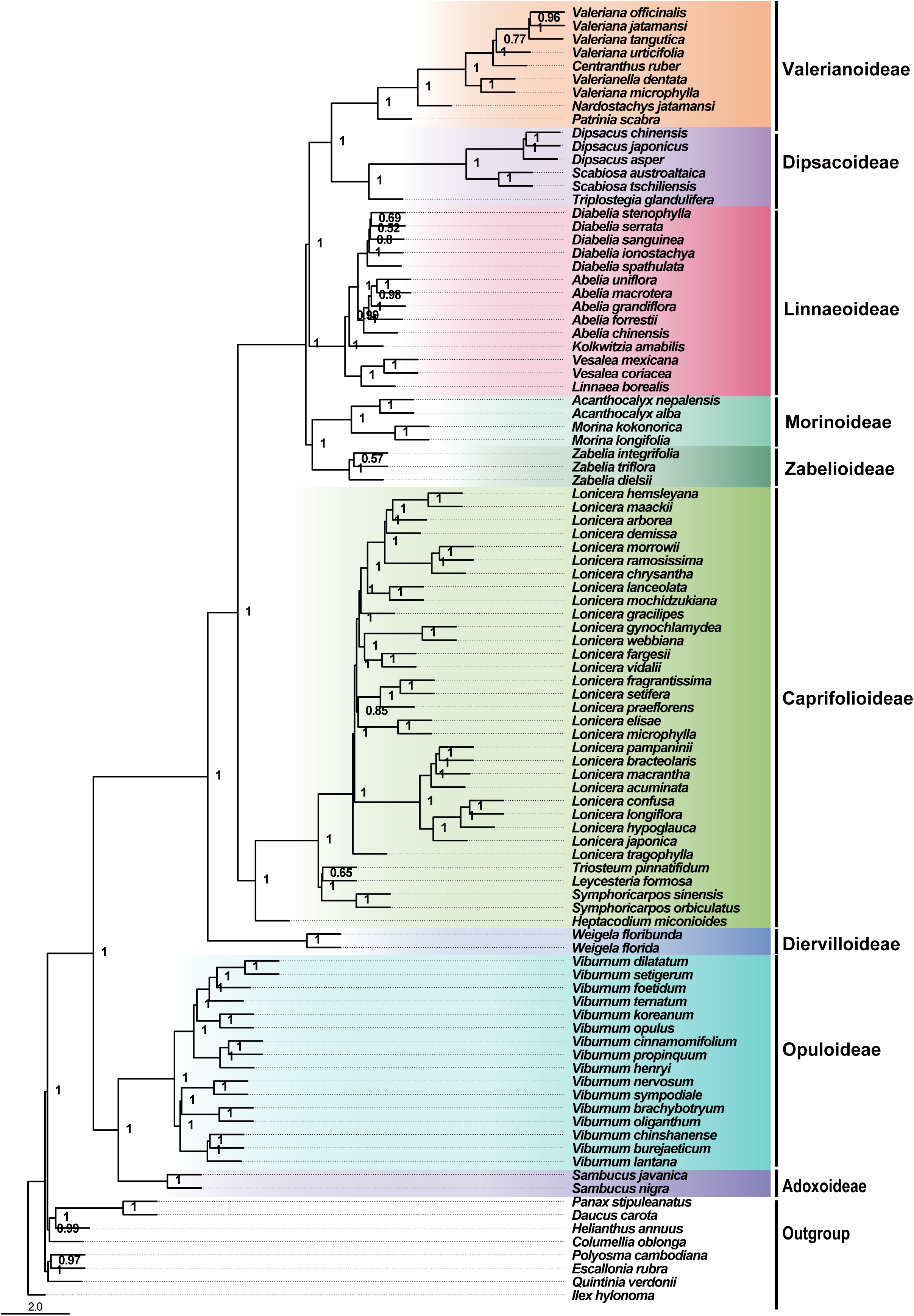
ASTRAL species tree of Dipsacales based on first and second codon positions (NT12) derived from Dataset 1. The NT12 data retained only the first and second codon positions. Node labels show local posterior probabilities (LPPs). Background shading denotes subfamilies. Branch lengths are measured in coalescent units.

**Figure S7.**
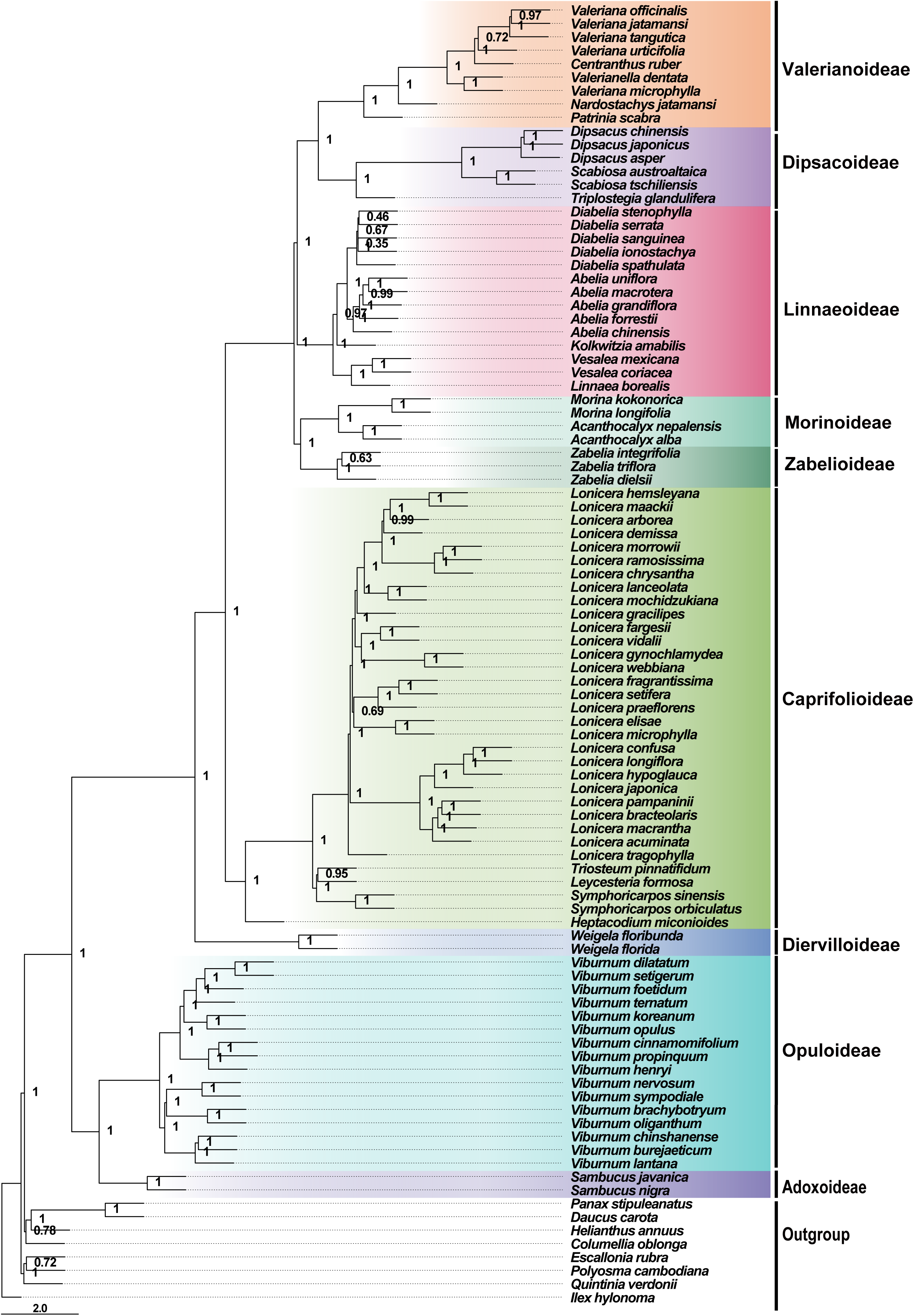
ASTRAL species tree of Dipsacales based on degeneracy-coded first and second codon positions (NT12-deg) derived from Dataset 1. The NT12-deg data were generated by degeneracy coding the first and second codon positions. Node labels show local posterior probabilities (LPPs). Background shading denotes subfamilies. Branch lengths are measured in coalescent units.

**Figure S8.**
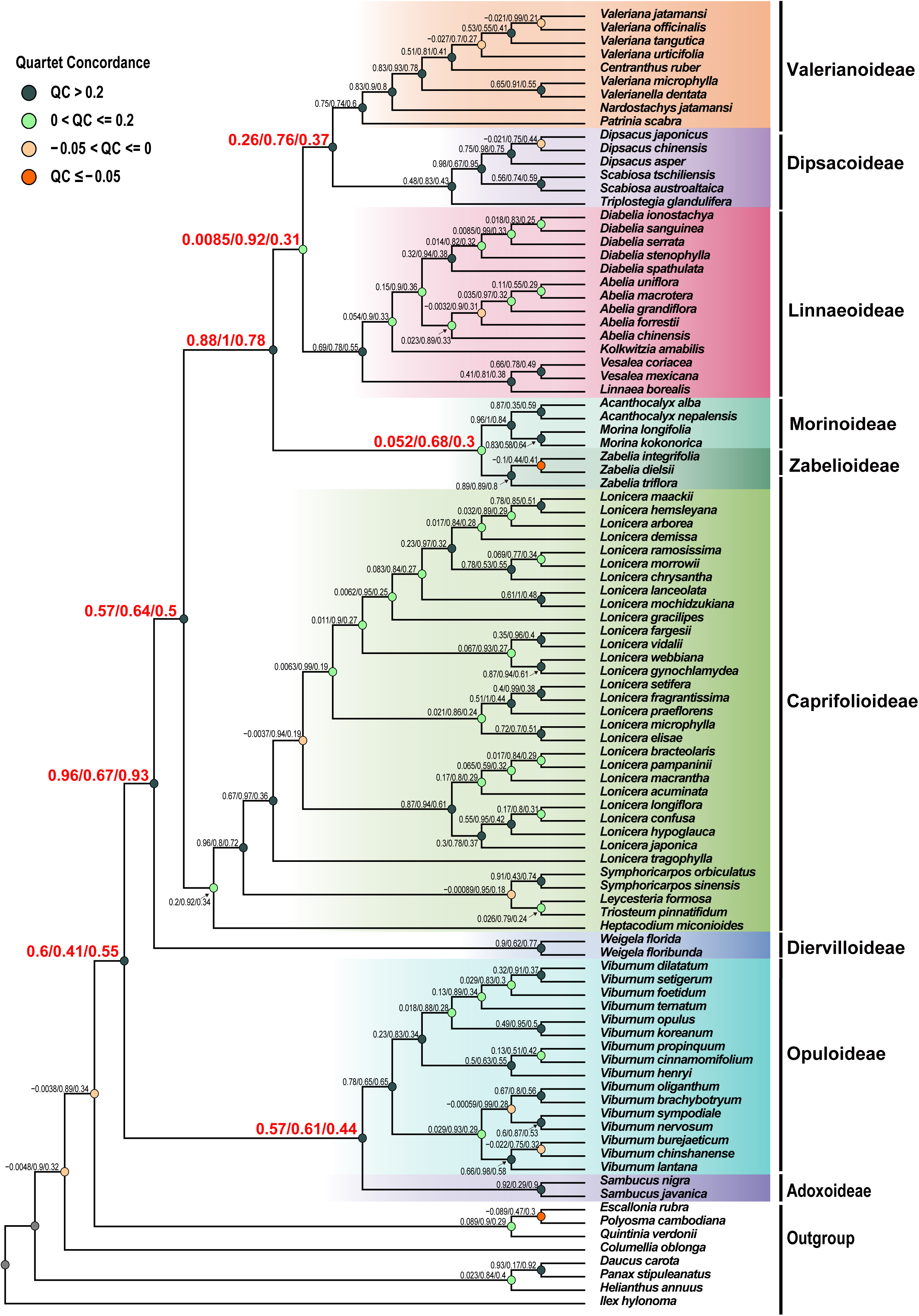
Quartet Sampling support across all internal nodes of the Dataset 1 species tree. Values at nodes indicate QC, QD, and QI, respectively. Circle colors represent different ranges of QC values, and red labels highlight the major inter-subfamily backbone nodes. Background colors identify the subfamilies.

**Figure S9.**
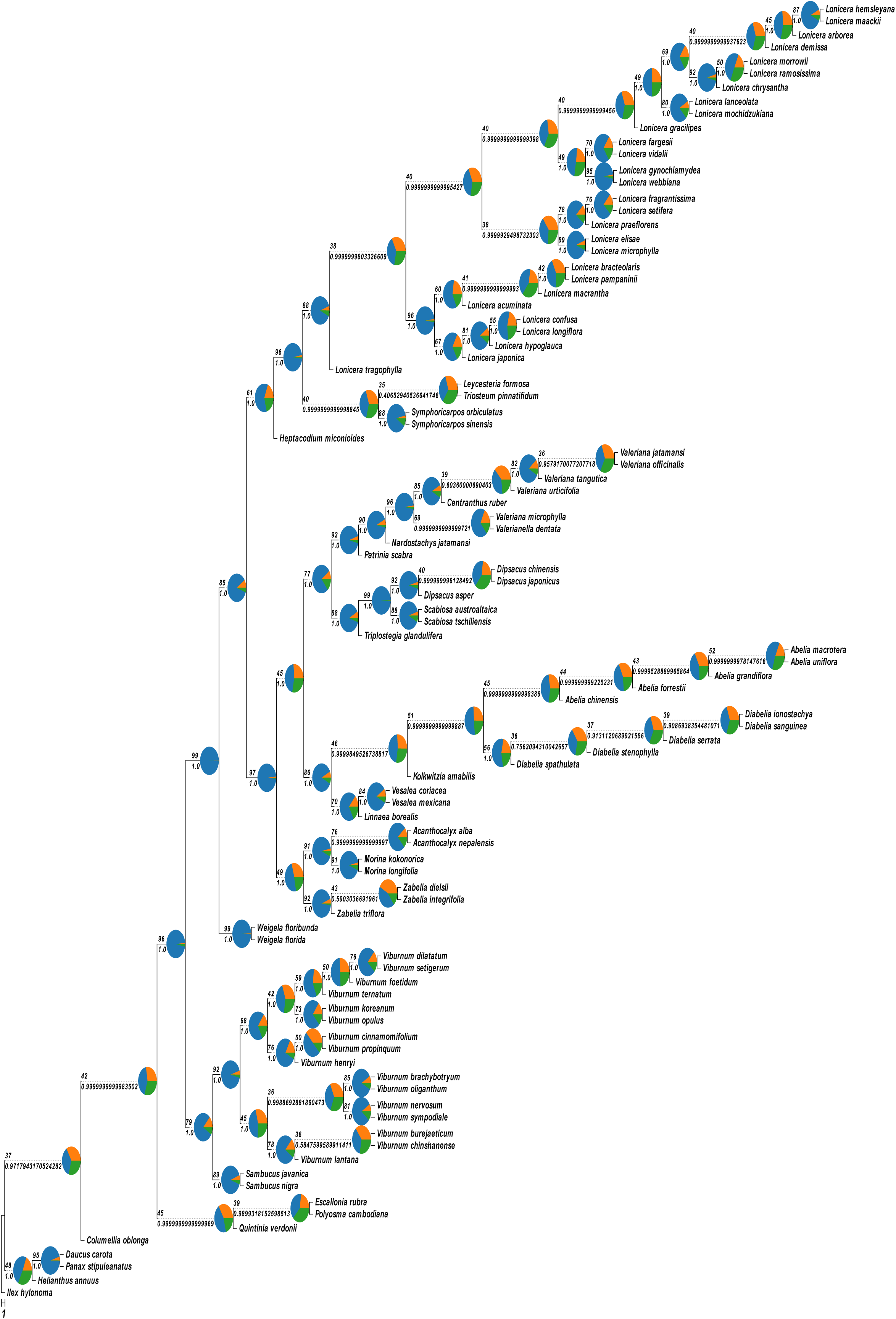
Gene-tree topology proportions across all internal nodes of the Dipsacales species tree. Pie charts show the proportions of gene trees supporting the species-tree topology and the two alternative topologies, illustrating the distribution of gene-tree concordance and conflict across the full tree.

**Figure S10.**
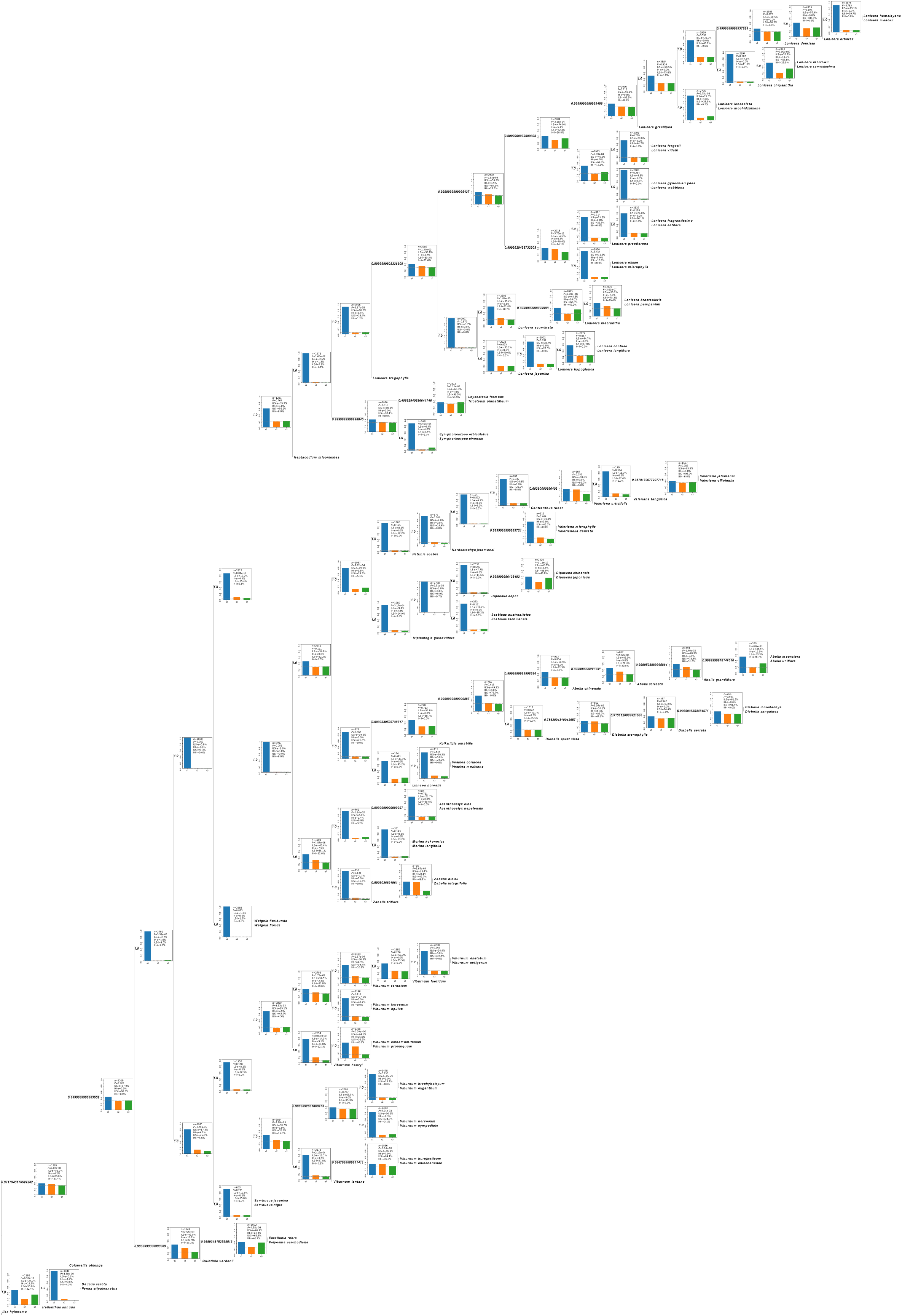
Gene-tree topology frequencies across all internal nodes of the Dipsacales species tree. Bar plots show the frequencies of gene trees supporting the main topology and the two alternative topologies, allowing comparison of gene-tree discordance among internal nodes.

## Tables

Table S1. Taxon sampling, classification, collection locality, voucher information, and SRA accession numbers for the 97 taxa included in this study.

Table S2. Publicly available transcriptome and genome datasets used to construct Dipsacales-specific nuclear reference locus sets

Table S3. Summary statistics of the two nuclear phylogenomic datasets

Table S4. Ranking of candidate taxon combinations for Dataset 2. Combinations were ranked by shared locus count, followed by lower mean missingness and greater total effective sequence length. The highest-ranking combination was selected for Dataset 2.

## Notes

### Competing Interest Statement

The authors have declared no competing interest.

### Funding

- National Natural Science Foundation of China: 32670274
- National Natural Science Foundation of China: 32270221

