## Supplementary material for "Nuclear phylogenomics illuminates deep lineage divergence and complex evolutionary histories in Dipsacales": Fig. S1.pdf

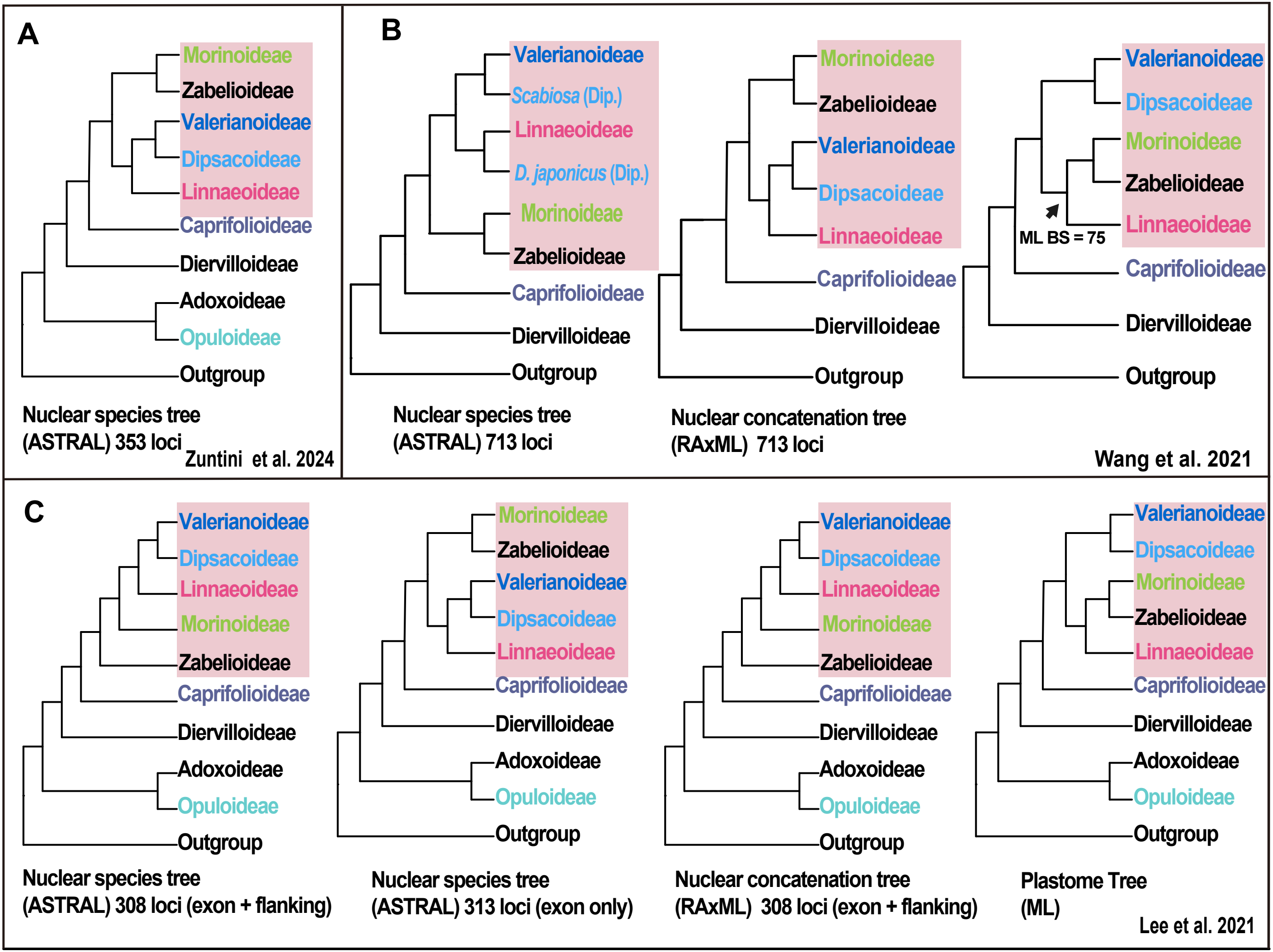
A
Morinoideae
Zabelioideae
Valerianoideae
Dipsacoideae
Linnaeoideae
Caprifolioideae
Diervilloideae
Adoxoideae
Opuloideae
Outgroup
Nuclear species tree
(ASTRAL) 353 loci
Zuntini et al. 2024
B
Valerianoideae
Scabiosa (Dip.)
Linnaeoideae
D. japonicus (Dip.)
Morinoideae
Zabelioideae
Caprifolioideae
Diervilloideae
Outgroup
Nuclear species tree
(ASTRAL) 713 loci
Morinoideae
Zabelioideae
Valerianoideae
Dipsacoideae
Linnaeoideae
Caprifolioideae
Diervilloideae
Outgroup
Nuclear concatenation tree
(RAxML) 713 loci
Valerianoideae
Dipsacoideae
Morinoideae
Zabelioideae
Linnaeoideae
ML BS = 75
Caprifolioideae
Diervilloideae
Outgroup
Wang et al. 2021
C
Valerianoideae
Dipsacoideae
Linnaeoideae
Morinoideae
Zabelioideae
Caprifolioideae
Diervilloideae
Adoxoideae
Opuloideae
Outgroup
Nuclear species tree
(ASTRAL) 308 loci (exon + flanking)
Morinoideae
Zabelioideae
Valerianoideae
Dipsacoideae
Linnaeoideae
Caprifolioideae
Diervilloideae
Adoxoideae
Opuloideae
Outgroup
Nuclear species tree
(ASTRAL) 313 loci (exon only)
Valerianoideae
Dipsacoideae
Linnaeoideae
Morinoideae
Zabelioideae
Caprifolioideae
Diervilloideae
Adoxoideae
Opuloideae
Outgroup
Nuclear concatenation tree
(RAxML) 308 loci (exon + flanking)
Valerianoideae
Dipsacoideae
Morinoideae
Zabelioideae
Linnaeoideae
Caprifolioideae
Diervilloideae
Adoxoideae
Opuloideae
Outgroup
Plastome Tree
(ML)
Lee et al. 2021
