## Supplementary material for "Nuclear phylogenomics illuminates deep lineage divergence and complex evolutionary histories in Dipsacales": Fig. S2.pdf

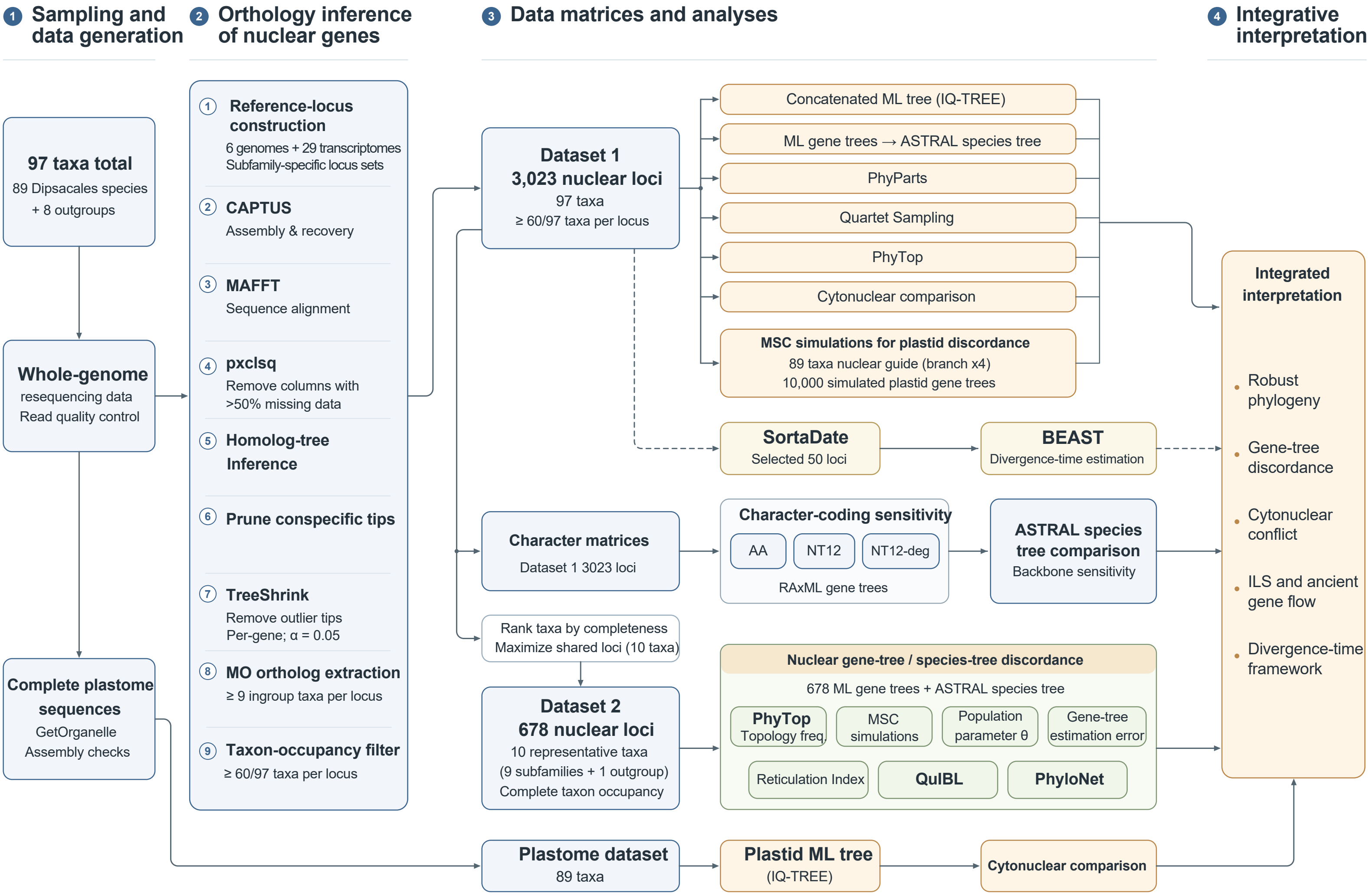

1 Sampling and data generation
2 Orthology inference of nuclear genes
3 Data matrices and analyses
4 Integrative interpretation
97 taxa total
89 Dipsacales species
+ 8 outgroups
Whole-genome resequencing data
Read quality control
Complete plastome sequences
GetOrganelle
Assembly checks
1 Reference-locus construction
6 genomes + 29 transcriptomes
Subfamily-specific locus sets
2 CAPTUS
Assembly & recovery
3 MAFFT
Sequence alignment
4 pxclsq
Remove columns with >50% missing data
5 Homolog-tree Inference
6 Prune conspecific tips
7 TreeShrink
Remove outlier tips
Per-gene; α = 0.05
8 MO ortholog extraction
≥ 9 ingroup taxa per locus
9 Taxon-occupancy filter
≥ 60/97 taxa per locus
Dataset 1
3,023 nuclear loci
97 taxa
≥ 60/97 taxa per locus
Concatenated ML tree (IQ-TREE)
ML gene trees → ASTRAL species tree
PhyParts
Quartet Sampling
PhyTop
Cytonuclear comparison
MSC simulations for plastid discordance
89 taxa nuclear guide (branch x4)
10,000 simulated plastid gene trees
SortaDate
Selected 50 loci
BEAST
Divergence-time estimation
Character matrices
Dataset 1 3023 loci
Character-coding sensitivity
AA
NT12
NT12-deg
RAxML gene trees
ASTRAL species tree comparison
Backbone sensitivity
Rank taxa by completeness
Maximize shared loci (10 taxa)
Dataset 2
678 nuclear loci
10 representative taxa
(9 subfamilies + 1 outgroup)
Complete taxon occupancy
Nuclear gene-tree / species-tree discordance
678 ML gene trees + ASTRAL species tree
PhyTop
Topology freq.
MSC simulations
Population parameter θ
Gene-tree estimation error
Reticulation Index
QuIBL
PhyloNet
Plastome dataset
89 taxa
Plastid ML tree
(IQ-TREE)
Cytonuclear comparison
Integrated interpretation
Robust phylogeny
Gene-tree discordance
Cytonuclear conflict
ILS and ancient gene flow
Divergence-time framework
