## Supplementary material for "Nuclear phylogenomics illuminates deep lineage divergence and complex evolutionary histories in Dipsacales": Fig. S4.pdf

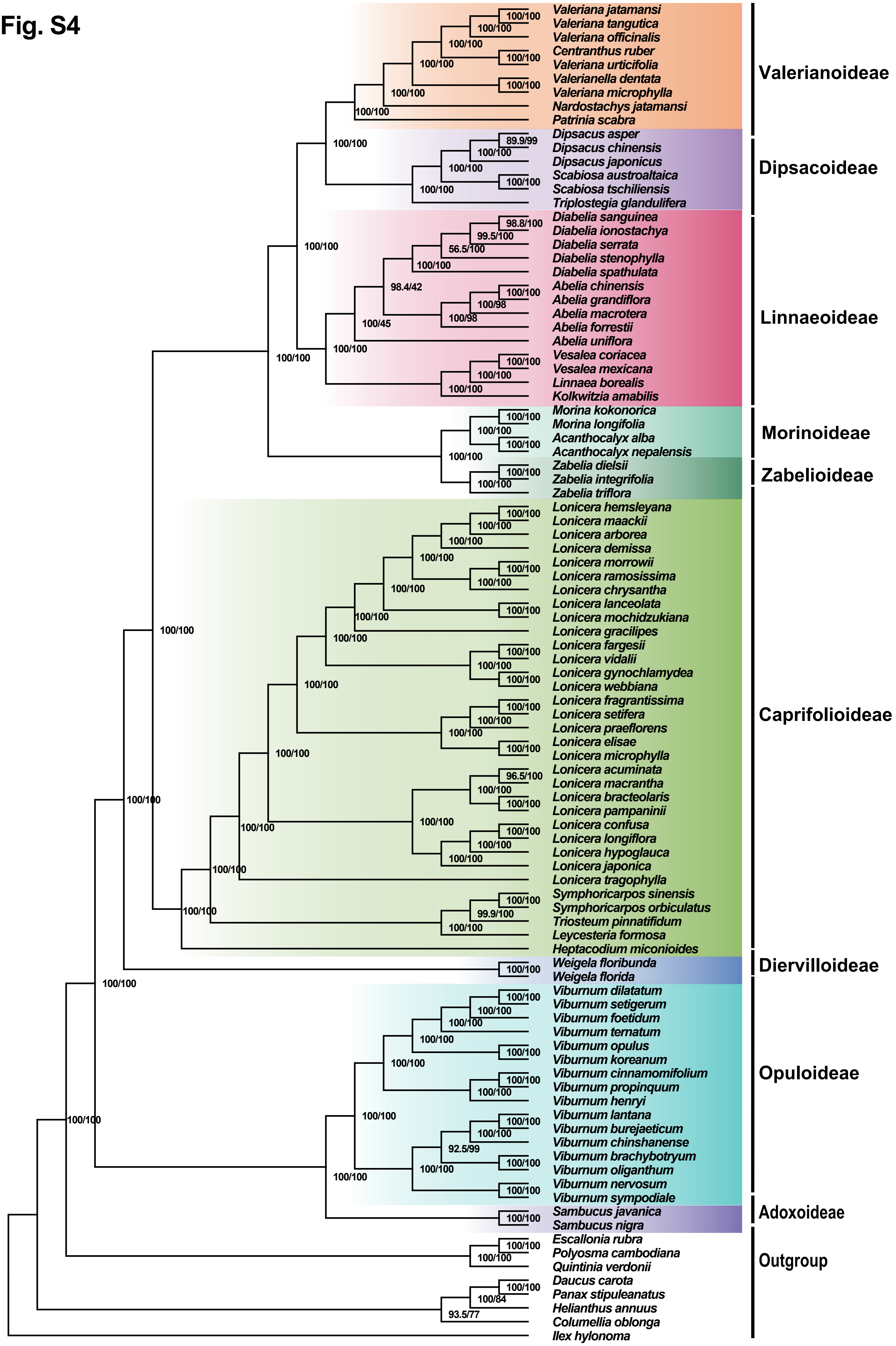
Valeriana jatamansi
Valeriana tangutica
Valeriana officinalis
Centranthus ruber
Valeriana urticifolia
Valerianella dentata
Valeriana microphylla
Nardostachys jatamansi
Patrinia scabra
Dipsacus asper
Dipsacus chinensis
Dipsacus japonicus
Scabiosa austroaltaica
Scabiosa tschiliensis
Triplostegia glandulifera
Diabelia sanguinea
Diabelia ionostachya
Diabelia serrata
Diabelia stenophylla
Diabelia spathulata
Abelia chinensis
Abelia grandiflora
Abelia macrotera
Abelia forrestii
Abelia uniflora
Vesalea coriacea
Vesalea mexicana
Linnaea borealis
Kolkwitzia amabilis
Morina kokonorica
Morina longifolia
Acanthocalyx alba
Acanthocalyx nepalensis
Zabelia dielsii
Zabelia integrifolia
Zabelia triflora
Lonicera hemsleyana
Lonicera maackii
Lonicera arborea
Lonicera demissa
Lonicera morrowii
Lonicera ramosissima
Lonicera chrysantha
Lonicera lanceolata
Lonicera mochidzukiana
Lonicera gracilipes
Lonicera fargesii
Lonicera vidalii
Lonicera gynochlamydea
Lonicera webbiana
Lonicera fragrantissima
Lonicera setifera
Lonicera praeflorens
Lonicera elisae
Lonicera microphylla
Lonicera acuminata
Lonicera macrantha
Lonicera bracteolaris
Lonicera pampaninii
Lonicera confusa
Lonicera longiflora
Lonicera hypoglauca
Lonicera japonica
Lonicera tragophylla
Symphoricarpos sinensis
Symphoricarpos orbiculatus
Triosteum pinnatifidum
Leycesteria formosa
Heptacodium miconioides
Weigela floribunda
Weigela florida
Viburnum dilatatum
Viburnum setigerum
Viburnum foetidum
Viburnum ternatum
Viburnum opulus
Viburnum koreanum
Viburnum cinnamomifolium
Viburnum propinquum
Viburnum henryi
Viburnum lantana
Viburnum burejaeticum
Viburnum chinshanense
Viburnum brachybotryum
Viburnum oliganthum
Viburnum nervosum
Viburnum sympodiale
Sambucus javanica
Sambucus nigra
Escallonia rubra
Polyosma cambodiana
Quintinia verdonii
Daucus carota
Panax stipuleanatus
Helianthus annuus
Columellia oblonga
Ilex hylonoma
Valerianoideae
Dipsacoideae
Linnaeoideae
Morinoideae
Zabelioideae
Caprifolioideae
Diervilloideae
Opuloideae
Adoxoideae
Outgroup
89.9/99
98.8/100
99.5/100
56.5/100
98.4/42
100/45
100/98
96.5/100
99.9/100
92.5/99
100/84
93.5/77
100/100
