## Supplementary material for "Nuclear phylogenomics illuminates deep lineage divergence and complex evolutionary histories in Dipsacales": Fig. S7.pdf

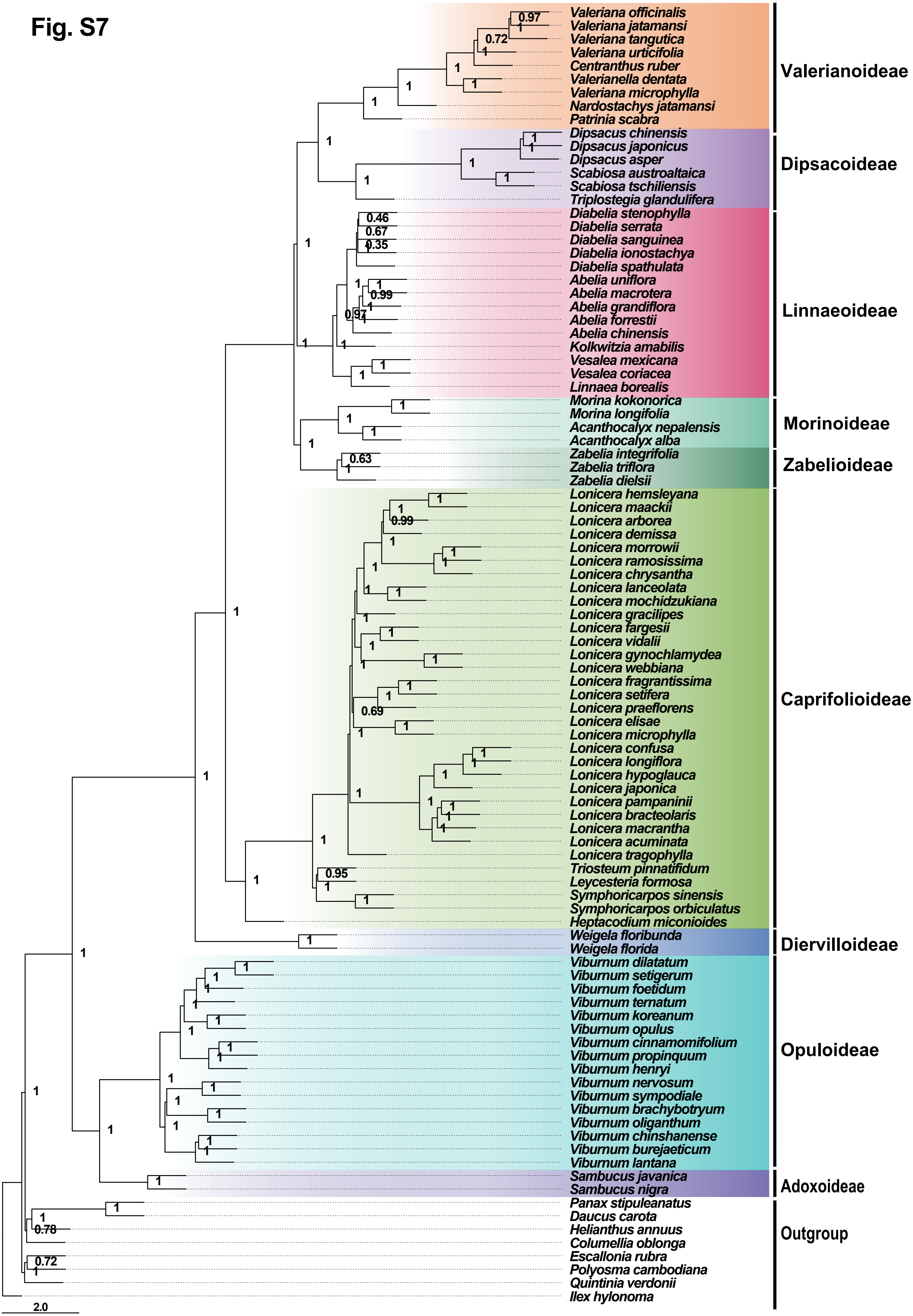
Valeriana officinalis
Valeriana jatamansi
Valeriana tangutica
Valeriana urticifolia
Centranthus ruber
Valerianella dentata
Valeriana microphylla
Nardostachys jatamansi
Patrinia scabra
Valerianoideae
Dipsacus chinensis
Dipsacus japonicus
Dipsacus asper
Scabiosa austroaltaica
Scabiosa tschiliensis
Triplostegia glandulifera
Dipsacoideae
Diabelia stenophylla
Diabelia serrata
Diabelia sanguinea
Diabelia ionostachya
Diabelia spathulata
Abelia uniflora
Abelia macrotera
Abelia grandiflora
Abelia forrestii
Abelia chinensis
Kolkwitzia amabilis
Vesalea mexicana
Vesalea coriacea
Linnaea borealis
Linnaeoideae
Morina kokonorica
Morina longifolia
Acanthocalyx nepalensis
Acanthocalyx alba
Morinoideae
Zabelia integrifolia
Zabelia triflora
Zabelia dielsii
Zabelioideae
Lonicera hemsleyana
Lonicera maackii
Lonicera arborea
Lonicera demissa
Lonicera morrowii
Lonicera ramosissima
Lonicera chrysantha
Lonicera lanceolata
Lonicera mochidzukiana
Lonicera gracilipes
Lonicera fargesii
Lonicera vidalii
Lonicera gynochlamydea
Lonicera webbiana
Lonicera fragrantissima
Lonicera setifera
Lonicera praeflorens
Lonicera elisae
Lonicera microphylla
Lonicera confusa
Lonicera longiflora
Lonicera hypoglauca
Lonicera japonica
Lonicera pampaninii
Lonicera bracteolaris
Lonicera macrantha
Lonicera acuminata
Lonicera tragophylla
Triosteum pinnatifidum
Leycesteria formosa
Symphoricarpos sinensis
Symphoricarpos orbiculatus
Heptacodium miconioides
Caprifolioideae
Weigela floribunda
Weigela florida
Diervilloideae
Viburnum dilatatum
Viburnum setigerum
Viburnum foetidum
Viburnum ternatum
Viburnum koreanum
Viburnum opulus
Viburnum cinnamomifolium
Viburnum propinquum
Viburnum henryi
Viburnum nervosum
Viburnum sympodiale
Viburnum brachybotryum
Viburnum oliganthum
Viburnum chinshanense
Viburnum burejaeticum
Viburnum lantana
Opuloideae
Sambucus javanica
Sambucus nigra
Adoxoideae
Panax stipuleanatus
Daucus carota
Helianthus annuus
Columellia oblonga
Escallonia rubra
Polyosma cambodiana
Quintinia verdonii
Ilex hylonoma
Outgroup
2.0
