## Supplementary material for "Nuclear phylogenomics illuminates deep lineage divergence and complex evolutionary histories in Dipsacales": Fig. S8.pdf

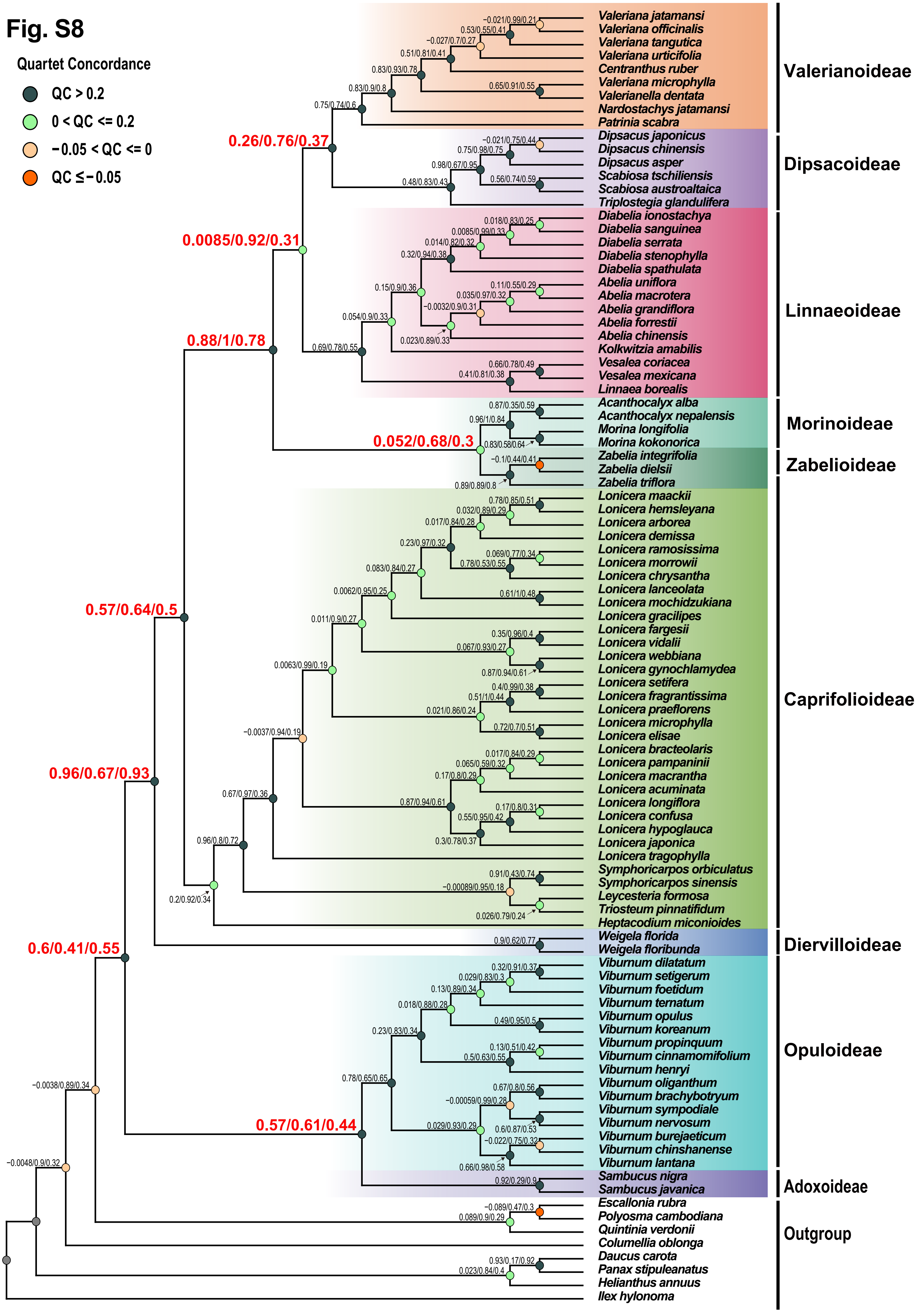
Quartet Concordance
QC > 0.2
0 < QC <= 0.2
−0.05 < QC <= 0
QC ≤ −0.05
0.26/0.76/0.37
0.0085/0.92/0.31
0.88/1/0.78
0.052/0.68/0.3
0.57/0.64/0.5
0.96/0.67/0.93
0.6/0.41/0.55
0.57/0.61/0.44
Valeriana jatamansi
Valeriana officinalis
Valeriana tangutica
Valeriana urticifolia
Centranthus ruber
Valeriana microphylla
Valerianella dentata
Nardostachys jatamansi
Patrinia scabra
Dipsacus japonicus
Dipsacus chinensis
Dipsacus asper
Scabiosa tschiliensis
Scabiosa austroaltaica
Triplostegia glandulifera
Diabelia ionostachya
Diabelia sanguinea
Diabelia serrata
Diabelia stenophylla
Diabelia spathulata
Abelia uniflora
Abelia macrotera
Abelia grandiflora
Abelia forrestii
Abelia chinensis
Kolkwitzia amabilis
Vesalea coriacea
Vesalea mexicana
Linnaea borealis
Acanthocalyx alba
Acanthocalyx nepalensis
Morina longifolia
Morina kokonorica
Zabelia integrifolia
Zabelia dielsii
Zabelia triflora
Lonicera maackii
Lonicera hemsleyana
Lonicera arborea
Lonicera demissa
Lonicera ramosissima
Lonicera morrowii
Lonicera chrysantha
Lonicera lanceolata
Lonicera mochidzukiana
Lonicera gracilipes
Lonicera fargesii
Lonicera vidalii
Lonicera webbiana
Lonicera gynochlamydea
Lonicera setifera
Lonicera fragrantissima
Lonicera praeflorens
Lonicera microphylla
Lonicera elisae
Lonicera bracteolaris
Lonicera pampaninii
Lonicera macrantha
Lonicera acuminata
Lonicera longiflora
Lonicera confusa
Lonicera hypoglauca
Lonicera japonica
Lonicera tragophylla
Symphoricarpos orbiculatus
Symphoricarpos sinensis
Leycesteria formosa
Triosteum pinnatifidum
Heptacodium miconioides
Weigela florida
Weigela floribunda
Viburnum dilatatum
Viburnum setigerum
Viburnum foetidum
Viburnum ternatum
Viburnum opulus
Viburnum koreanum
Viburnum propinquum
Viburnum cinnamomifolium
Viburnum henryi
Viburnum oliganthum
Viburnum brachybotryum
Viburnum sympodiale
Viburnum nervosum
Viburnum burejaeticum
Viburnum chinshanense
Viburnum lantana
Sambucus nigra
Sambucus javanica
Escallonia rubra
Polyosma cambodiana
Quintinia verdonii
Columellia oblonga
Daucus carota
Panax stipuleanatus
Helianthus annuus
Ilex hylonoma
Valerianoideae
Dipsacoideae
Linnaeoideae
Morinoideae
Zabelioideae
Caprifolioideae
Diervilloideae
Opuloideae
Adoxoideae
Outgroup
