## Supplementary material for "Nuclear phylogenomics illuminates deep lineage divergence and complex evolutionary histories in Dipsacales": Fig. S9.pdf

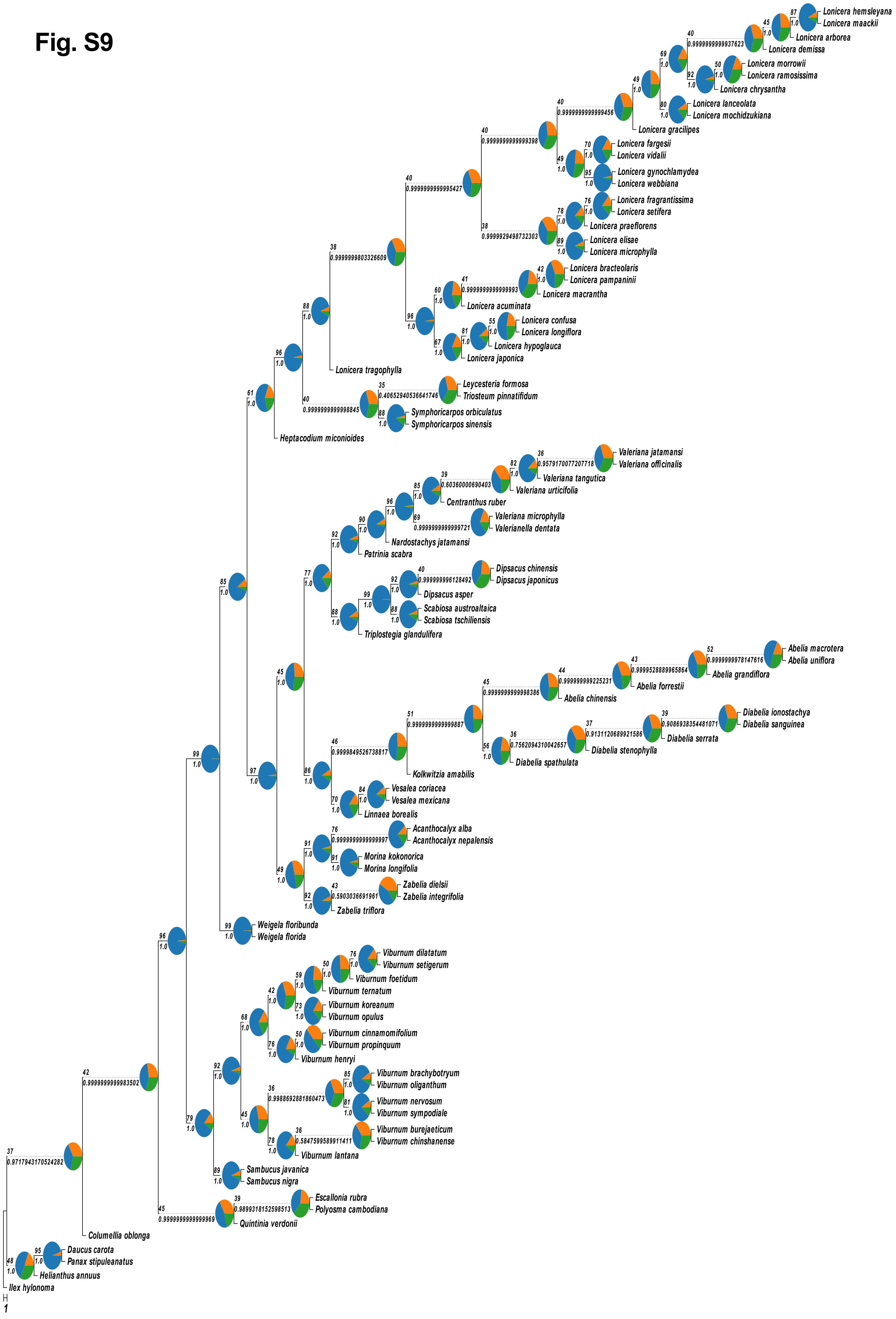
87
1.0
Lonicera hemsleyana
Lonicera maackii
45
1.0
Lonicera arborea
40
0.9999999999937623
Lonicera demissa
69
1.0
50
1.0
Lonicera morrowii
Lonicera ramosissima
92
1.0
Lonicera chrysantha
49
1.0
80
1.0
Lonicera lanceolata
Lonicera mochidzukiana
40
0.9999999999999456
Lonicera gracilipes
40
0.9999999999999398
70
1.0
Lonicera fargesii
Lonicera vidalii
49
1.0
95
1.0
Lonicera gynochlamydea
Lonicera webbiana
40
0.9999999999995427
76
1.0
Lonicera fragrantissima
Lonicera setifera
78
1.0
Lonicera praeflorens
38
0.9999929498732303
89
1.0
Lonicera elisae
Lonicera microphylla
38
0.9999999803326609
42
1.0
Lonicera bracteolaris
Lonicera pampaninii
41
0.9999999999999993
Lonicera macrantha
60
1.0
Lonicera acuminata
55
1.0
Lonicera confusa
Lonicera longiflora
96
1.0
81
1.0
Lonicera hypoglauca
67
1.0
Lonicera japonica
88
1.0
96
1.0
Lonicera tragophylla
35
0.406529405366641746
Leycesteria formosa
Triosteum pinnatifidum
40
0.9999999999998845
88
1.0
Symphoricarpos orbiculatus
Symphoricarpos sinensis
61
1.0
Heptacodium miconioides
36
0.9579170077207718
Valeriana jatamansi
Valeriana officinalis
82
1.0
Valeriana tangutica
39
0.60360000690403
Valeriana urticifolia
85
1.0
Centranthus ruber
96
1.0
69
0.9999999999999721
Valeriana microphylla
Valerianella dentata
90
1.0
Nardostachys jatamansi
92
1.0
Patrinia scabra
40
0.999999996128492
Dipsacus chinensis
Dipsacus japonicus
92
1.0
Dipsacus asper
99
1.0
88
1.0
Scabiosa austroaltaica
Scabiosa tschiliensis
77
1.0
88
1.0
Triplostegia glandulifera
85
1.0
52
0.9999999978147616
Abelia macrotera
Abelia uniflora
43
0.9999528889965864
Abelia grandiflora
44
0.999999999225231
Abelia forrestii
45
0.9999999999998386
Abelia chinensis
39
0.9086938354481071
Diabelia ionostachya
Diabelia sanguinea
37
0.9131120689921586
Diabelia serrata
51
0.9999999999999887
36
0.7562094310042657
Diabelia stenophylla
56
1.0
Diabelia spathulata
46
0.9999849526738817
Kolkwitzia amabilis
84
1.0
Vesalea coriacea
Vesalea mexicana
86
1.0
70
1.0
Linnaea borealis
45
1.0
97
1.0
76
0.9999999999999997
Acanthocalyx alba
Acanthocalyx nepalensis
91
1.0
91
1.0
Morina kokonorica
Morina longifolia
49
1.0
43
0.5903036691961
Zabelia dielsii
Zabelia integrifolia
92
1.0
Zabelia triflora
99
1.0
Weigela floribunda
Weigela florida
99
1.0
96
1.0
76
1.0
Viburnum dilatatum
Viburnum setigerum
50
1.0
Viburnum foetidum
59
1.0
Viburnum ternatum
42
1.0
73
1.0
Viburnum koreanum
Viburnum opulus
68
1.0
50
1.0
Viburnum cinnamomifolium
Viburnum propinquum
76
1.0
Viburnum henryi
92
1.0
85
1.0
Viburnum brachybotryum
Viburnum oliganthum
36
0.9988692881860473
81
1.0
Viburnum nervosum
Viburnum sympodiale
45
1.0
36
0.5847595589911411
Viburnum burejaeticum
Viburnum chinshanense
78
1.0
Viburnum lantana
79
1.0
89
1.0
Sambucus javanica
Sambucus nigra
42
0.9999999999983502
39
0.9899318152598513
Escallonia rubra
Polyosma cambodiana
45
0.9999999999999969
Quintinia verdonii
37
0.9717943170524282
Columellia oblonga
95
1.0
Daucus carota
Panax stipuleanatus
48
1.0
Helianthus annuus
Ilex hylonoma
1
