## Supplementary material for "Nuclear phylogenomics illuminates deep lineage divergence and complex evolutionary histories in Dipsacales": Fig. S10.pdf

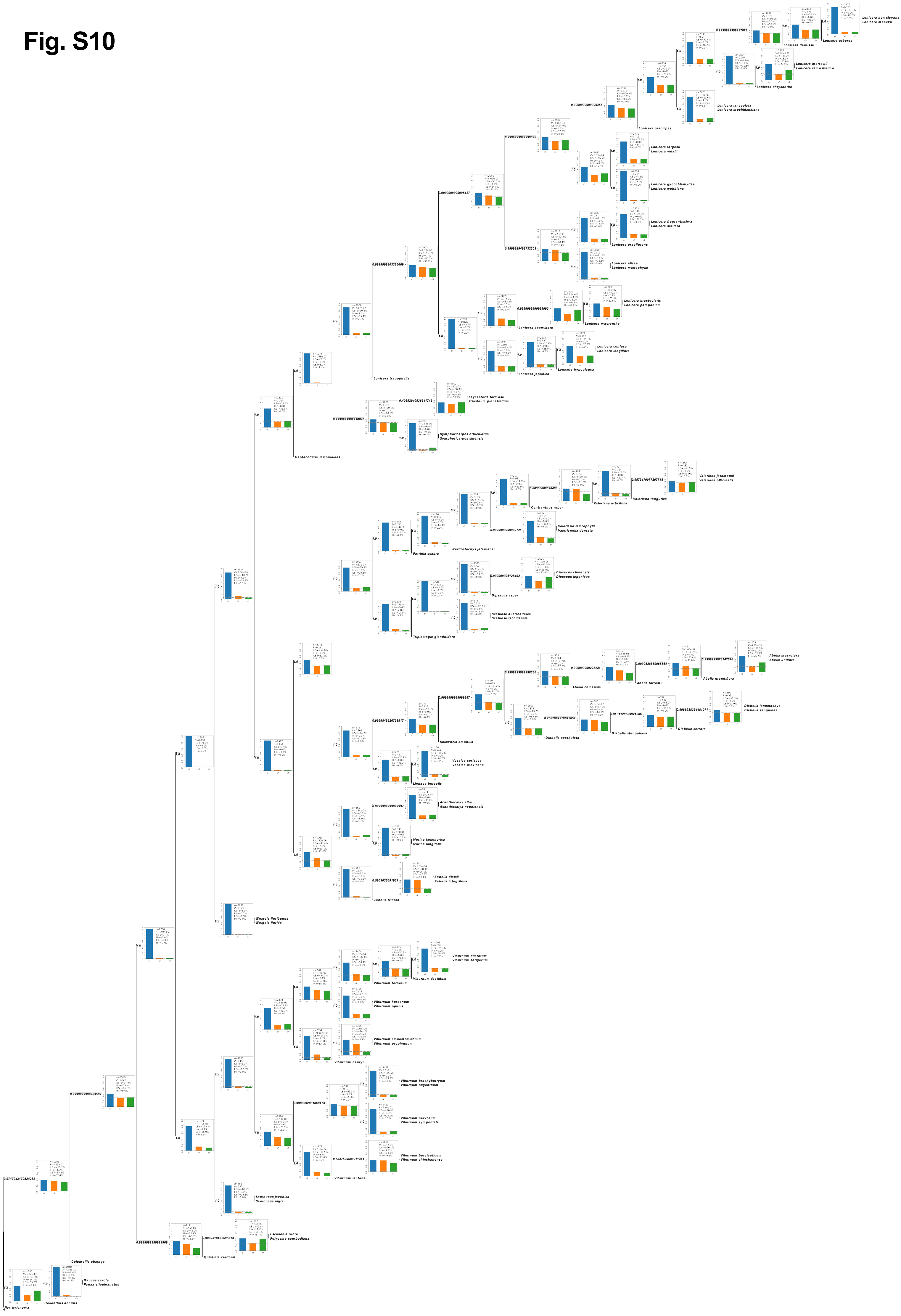

Lonicera hemsleyana
Lonicera maackii
Lonicera arborea
Lonicera demissa
Lonicera morrowii
Lonicera ramosissima
Lonicera chrysantha
Lonicera lanceolata
Lonicera mochidzukiana
Lonicera gracilipes
Lonicera fargesii
Lonicera vidalii
Lonicera gynochlamydea
Lonicera webbiana
Lonicera fragrantissima
Lonicera setifera
Lonicera praeflorens
Lonicera elisae
Lonicera microphylla
Lonicera bracteolaris
Lonicera pampaninii
Lonicera acuminata
Lonicera macrantha
Lonicera confusa
Lonicera longiflora
Lonicera japonica
Lonicera hypoglauca
Lonicera tragophylla
Leycesteria formosa
Triosteum pinnatifidum
Symphoricarpos orbiculatus
Symphoricarpos sinensis
Heptacodium miconioides
Valeriana jatamansi
Valeriana officinalis
Valeriana tangutica
Valeriana urticifolia
Centranthus ruber
Valeriana microphylla
Valerianella dentata
Nardostachys jatamansi
Patrinia scabra
Dipsacus chinensis
Dipsacus japonicus
Dipsacus asper
Scabiosa austroafrica
Scabiosa tschiliensis
Triplostegia glandulifera
Abelia macrotera
Abelia uniflora
Abelia grandiflora
Abelia forrestii
Abelia chinensis
Diabelia ionostachya
Diabelia sanguinea
Diabelia serrata
Diabelia stenophylla
Diabelia spathulata
Kolkwitzia amabilis
Vesalea coriacea
Vesalea mexicana
Linnaea borealis
Acanthocalyx alba
Acanthocalyx nepalensis
Morina kokonorica
Morina longifolia
Zabelia dielsii
Zabelia integrifolia
Zabelia triflora
Weigela floribunda
Weigela florida
Viburnum dilatatum
Viburnum setigerum
Viburnum foetidum
Viburnum ternatum
Viburnum koreanum
Viburnum opulus
Viburnum cinnamomifolium
Viburnum propinquum
Viburnum henryi
Viburnum brachybotryum
Viburnum oliganthum
Viburnum nervosum
Viburnum sympodiale
Viburnum burejaeticum
Viburnum chinshanense
Viburnum lantana
Sambucus javanica
Sambucus nigra
Escallonia rubra
Polyosma cambodiana
Quintinia verdonii
Columellia oblonga
Daucus carota
Panax stipuleanatus
Helianthus annuus
Ilex hylonoma
