## Supplementary figures and images for "Nuclear phylogenomics illuminates deep lineage divergence and complex evolutionary histories in Dipsacales"

### Fig. S3.pdf

Fig. S3

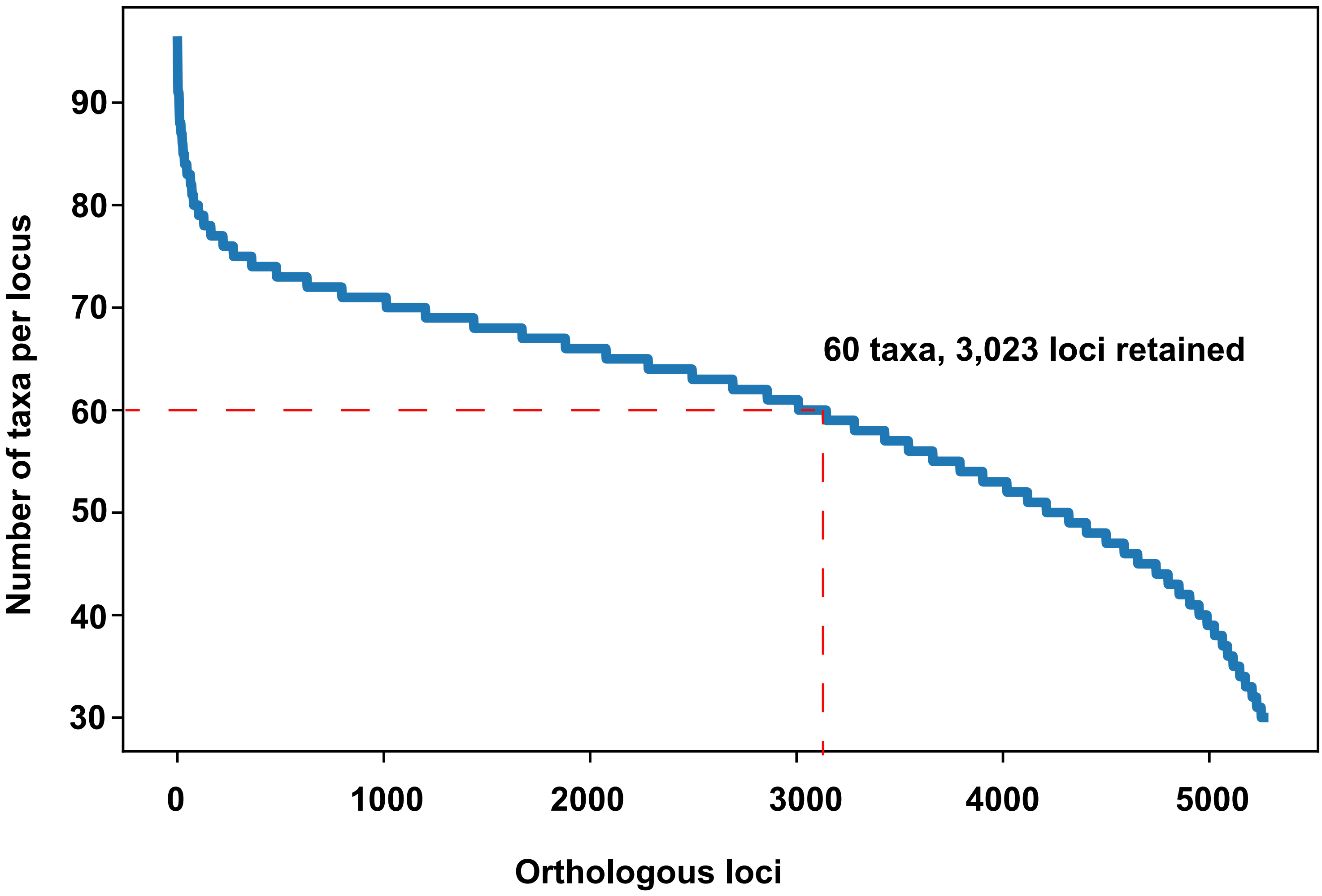

60 taxa, 3,023 loci retained
Number of taxa per locus
90
80
70
60
50
40
30
0
1000
2000
3000
4000
5000
Orthologous loci
